# “Population genomics reveals divergent lineages across Europe in the vulnerable mire plant *Drosera rotundifolia”*

**DOI:** 10.64898/2026.05.28.728421

**Authors:** Kristina Kuprina, Markus Sommer, Hildegard Kieninger, Malte Zörner, Martin Schnittler, Manuela Bog

## Abstract

Genomic variation within populations reflects both past and contemporary evolutionary processes. The genetic structure of the European peatland plant species is shaped by complex postglacial recolonization and recent habitat loss. Here, we investigate population genomic patterns in the vulnerable mire plant, round-leaved sundew (*Drosera rotundifolia* L.). Using ddRAD sequencing of 311 individuals from 38 populations across Europe, we detected significant genetic differentiation (Fst = 0.02-0.44, *p* < 0.05) and a consistent deficit of heterozygosity (Fis = 0.248) across populations. In contrast, sequencing of 10,449 bp of chloroplast DNA revealed extremely low variation, with only one SNP detected. Clustering analyses (Admixture and DAPC) both identified pronounced genetic structure comprising three major clusters (Western, Northern, and Eastern) that broadly correspond to European biogeographic regions. Genetic differentiation was partially explained by geographic distance (3.3%; *p* = 0.0001; Mantel test), while climatic variables, particularly temperature and precipitation, accounted for 2.9% of genomic variation (*p* < 0.0001; redundancy analysis). The observed pattern is consistent with polygenic responses to climatic gradients and is supported by 1,022 SNPs, distributed across 632 loci, that were significantly associated with environmental variables. Demographic reconstruction revealed distinct evolutionary trajectories among clusters, with the Western cluster showing a more recent expansion. Together, our results suggest that the genetic structure of *D. rotundifolia* is a result of postglacial recolonization, geographic isolation, and climate influence. We highlight the importance of conserving genetic diversity of *D. rotundifolia* across all three clusters and accounting for potential local adaptation and population vulnerability in future conservation strategies.

## 1. Introduction

Evolutionary history is encoded in genomes and reflected in patterns of population genetic variation. These patterns capture signatures of range expansions and contractions, barriers to gene flow, and adaptive processes (Hewitt 2000; Ellegren and Galtier 2016; Suda *et al*. 2025). Interpreting these signals allows us to detect demographic events such as bottlenecks following range contraction or founder effects following invasion or postglacial recolonization.

During the Last Glacial Maximum (LGM, ∼26,000–19,000 years ago), many European plant species survived in glacial refugia, areas with locally milder conditions where populations could persist despite ice-age climates (Heusser 1955). The main glacial refugia were located in southern Europe, particularly the Iberian, Italian, and Balkan peninsulas (Bennett *et al*. 1991). However, fossil and genetic data indicate that various species also survived farther north in smaller, scattered areas, such as the Carpathians, the Baltics, and in the Scandinavian microrefugia (Birks 2003; Willis and van Andel 2004; Stewart *et al*. 2010; Quinzin *et al*. 2017; Westergaard *et al*. 2019; Hošek *et al*. 2024). Consequently, the modern genetic structure of European plant species was shaped not by recolonization from a single uniform source, but by expansion from multiple, geographically distinct refugial populations.

The historical signals are also shaped by contemporary population characteristics, including size and connectivity. Small, isolated populations may differentiate substantially through genetic drift alone, even in the absence of adaptive pressure (Knowles and Richards 2005). The populations of wetland plants are especially affected by human activities like agriculture, urban sprawl and land reclamation. In Germany, for example, more than 95% of peatlands have experienced degradation due to drainage, primarily serving agricultural purposes such as crop culture, meadows or pastures (Abel *et al*. 2019; Tanneberger *et al*. 2021). Wetlands are also shown to be highly susceptible to climate change (Johnson and Poiani 2016; Antala *et al*. 2022). As a result, affected populations may become patchy and genetically isolated, with genetic structure shaped by both their evolutionary history and contemporary demography.

*Drosera rotundifolia* L. (round-leaved sundew) offers a compelling model system in which to examine these dynamics. This carnivorous plant has a circumboreal distribution, occurring in wetlands across Europe, North America, and Asia (Meusel *et al*. 1965; Wolf *et al*. 2006; Figure 1a). Beyond its core range, it also occurs in isolated populations in temperate and subtropical regions, with only two confirmed tropical populations (Philippines and Papua New Guinea; Fleischmann 2021). Even though this species has the largest global range of all *Drosera*, its habitat is strictly limited to nutrient-poor and mostly *Sphagnum*-dominated raised bogs (Baranyai and Joosten 2016), which have experienced extensive historical fragmentation during Pleistocene glaciation and continue to face contemporary threats from land-use change and climate-driven habitat loss (Fleischmann 2021; Fluet-Chouinard *et al*. 2023). Many extant populations of *D. rotundifolia* are small, isolated, and confined to discrete mire habitats, making them susceptible to genetic drift and reduced gene flow (Eschenbrenner *et al*. 2016; 2019).

**Figure 1.**
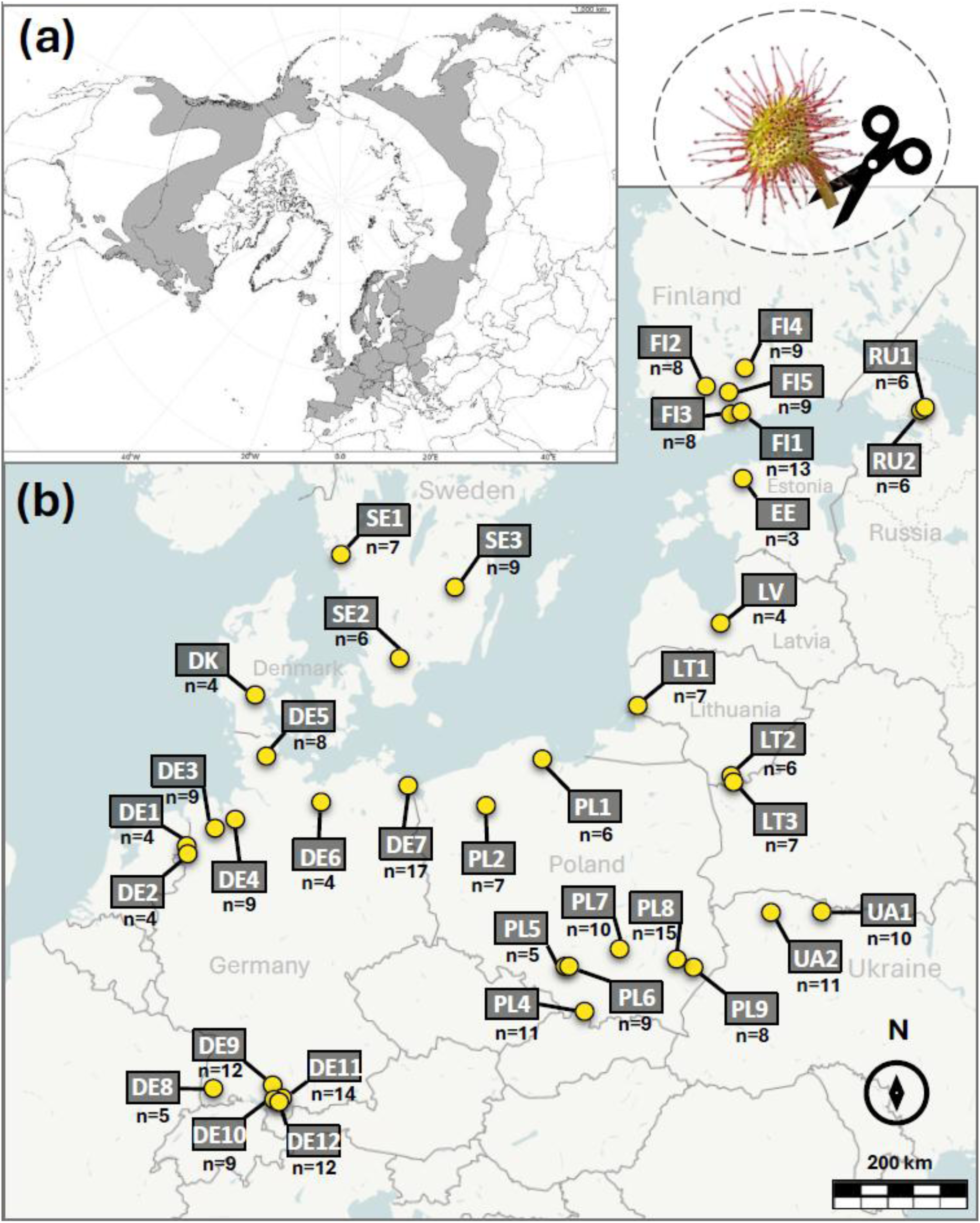
(a) Distribution of *Drosera rotundifolia* in Holoarctic (Baranyai and Joosten 2016); (b) Location of the 38 studied populations of *D. rotundifolia* across Europe. Numbers under population names indicate the sample size (n = 311 in total).

*D. rotundifolia* can reproduce both asexually and sexually. Asexual reproduction occurs by forming plantlets from the leaf buds or by forming a secondary rosette. In both cases, the distribution of clonal individuals is restricted locally. Sexual reproduction is considered to be the main reproductive pathway (Baranyai and Joosten 2016). The flowers of *D. rotundifolia* are hermaphroditic and predominantly autogamous (self-fertilizing) or cleistogamous, meaning flowers remain closed and self-pollinate (McPherson 2010). When flowers do open, they remain open for only a few hours, allowing limited insect pollination (McPherson 2010). Seed dispersal is restricted to only a few centimetres from the parent plant (Seeholzer 1993; Wolf *et al*. 2006; Baranyai and Joosten 2016). Additionally, unlike many plant species that are polyploid, *D. rotundifolia* has a diploid set of chromosomes (2n = 20; Kondo and Segawa 1988). Polyploidy can mask harmful recessive mutations (Soltis and Soltis 2000; Galloway *et al*. 2003; Rosche *et al*. 2017), whereas diploidy leaves such mutations exposed to selection by a higher inbreeding depression. Together with the habitat loss, we expect these characteristics to promote inbreeding and enhance genetic drift in *D. rotundifola*.

Despite its wide distribution, habitat specialization, and vulnerability to fragmentation, the population genetic structure and demographic history of *D. rotundifolia* remains poorly understood. To date, population genetic studies within the genus *Drosera* have been conducted exclusively on *D. rotundifolia*, with one study from Schleswig-Holstein, Germany (Eschenbrenner *et al*. 2019), and another from South Korea (Chung *et al*. 2013). These studies reported contrasting levels of within-population diversity: extremely low in North Korean populations (Chung *et al*. 2013; He = 0.005, allozymes) and relatively high in Schleswig-Holstein (Eschenbrenner *et al*. 2019; minimum He = 0.109, Inter-simple sequence repeats). These inconsistent patterns may reflect lower variation of allozyme marker and, as suggested by Eschenbrenner *et al*. (2019), lower number and size of glacial refugia that served as source populations for contemporary lineages. No publication addressed genetic structure of *D. rotundifolia* across its European range yet.

In this study, we address this gap by generating genome-wide population genetic data for *Drosera rotundifolia* from 38 European populations. Specifically, we characterize patterns of genetic diversity and population structure, infer demographic history and potential refugial sources, and evaluate priorities for conservation under ongoing habitat loss. We test the following hypotheses: (i) *D. rotundifolia* populations in Europe are genetically differentiated and exhibit high levels of inbreeding; (ii) large-scale patterns of genetic variation reflect the species’ biogeographic history; and (iii) both climatic variables and geographic distance contribute to shaping genetic structure.

## 2. Material and Methods

### 2.1. Plant material

Samples of *Drosera rotundifolia* were collected between June and September in 2023 and 2024. In total, 311 individuals from 38 populations were included in the study (Figure 1b). A population was defined as a group of individuals growing in a continuous peatland area not separated by natural or artificial barriers. Depending on the area size, we collected 3–17 individuals per population (Table 1): in small populations, sampling was quasi-random, whereas in larger peatlands, individuals were sampled along a transect. To reduce the likelihood of collecting clonal or closely related individuals, we maintained a minimum distance of 2 m between sampled plants. Each sample consisted of 1–2 leaves preserved in silica gel. Metadata (population information, geographic coordinates, data of collection, sequences names) for each sample are provided in Supplementary Table ST1.

**Table 1.** Population-wise and total estimates of genetic variation of *Drosera rotundifolia* populations. N - number of samples, MLGs – number of multilocus genotypes, C - clonal richness, Ho – adjusted observed heterozygosity, uHe – unbiased expected heterozygosity, Fis – inbreeding coefficient. SE – standard errors. Calculations of C, Ho, uHe and Fis were conducted only for populations with at least six MLGs.

| Populat<br>ion | Bog area,<br>~ ha | Maximum<br>distance<br>between<br>samples, m | N | MLGs | C | Ho | uHe | Fis |
| --- | --- | --- | --- | --- | --- | --- | --- | --- |
| DE1 | 409 | - | 4 | 1 | - | - | - | - |
| DE2 | 708 | 1689 | 4 | 3 | - | - | - | - |
| DE3 | 1.5 | 124 | 9 | 5 | 0.50 | 0.0113 | 0.0233 | 0.455 |
| DE4 | 230 | 148 | 9 | 5 | 0.50 | 0.0116 | 0.0262 | 0.493 |
| DE5 | 38 | 155 | 8 | 4 | - | - | - | - |
| DE6 | 567 | 1287 | 4 | 4 | - | - | - | - |
| DE7 | 0.5 | 148 | 17 | 7 | 0.44<sup>L</sup> | 0.0108 | 0.0180<sup>L</sup> | 0.337 |
| DE8 | 1 | 75 | 5 | 5 | 1.00<sup>H</sup> | 0.0116 | 0.0185 | 0.288<sup>L</sup> |
| DE9 | 1,812 | 1977 | 12 | 11 | 0.91 | 0.0101 | 0.0203 | 0.364 |
| DE10 | 40 | 440 | 9 | 3 | - | - | - | - |
| DE11 | 315 | 1156 | 14 | 12 | 0.79 | 0.0102 | 0.0248 | 0.438 |
| DE12 | 104 | 272 | 12 | 11 | 0.91 | 0.0133 | 0.0247 | 0.333 |
| DK | 12 | 31 | 4 | 2 | - | - | - | - |
| EE | 445 | 32 | 3 | 2 | - | - | - | - |
| FI1 | 10 | 1364 | 13 | 12 | 0.92 | 0.0156<sup>H</sup> | 0.0292<sup>H</sup> | 0.342 |
| FI2 | 3,041 | 2237 | 8 | 8 | 1.00<sup>H</sup> | 0.0109 | 0.0236 | 0.444 |
| FI3 | 285 | 2709 | 8 | 8 | 1.00<sup>H</sup> | 0.0115 | 0.0218 | 0.361 |
| FI4 | 121 | 2881 | 9 | 9 | 1.00<sup>H</sup> | 0.0118 | 0.0284 | 0.460 |
| FI5 | 3 | 448 | 9 | 6 | 0.63 | 0.0127 | 0.0217 | 0.349 |
| LT1 | 2,222 | - | 6 | 5 | 0.80 | 0.0121 | 0.0244 | 0.411 |
| LT2 | 1 | 107 | 7 | 2 | - | - | - | - |
| LT3 | 2 | 312 | 7 | 4 | - | - | - | - |
| LV | 364 | - | 4 | 2 | - | - | - | - |
| PL1 | 2 | 144 | 6 | 6 | 1.00<sup>H</sup> | 0.0105 | 0.0211 | 0.411 |
| PL2 | 7.5 | 75 | 7 | 5 | 0.67 | 0.0071<sup>L</sup> | 0.0193 | 0.547<sup>H</sup> |
| PL4 | 159 | 858 | 11 | 7 | 0.60 | 0.0092 | 0.0238 | 0.490 |
| PL5 | 27 | 578 | 5 | 3 | - | - | - | - |
| PL6 | 1.5 | 70 | 9 | 7 | 0.75 | 0.0138 | 0.0232 | 0.290 |
| PL7 | 33 | 483 | 10 | 7 | 0.67 | 0.0129 | 0.0239 | 0.388 |
| PL8 | 92 | 1567 | 15 | 13 | 0.86 | 0.0094 | 0.0214 | 0.410 |
| PL9 | 1 | 1086 | 8 | 6 | 0.71 | 0.0134 | 0.0253 | 0.393 |
| RU1 | 0.5 | 153 | 6 | 5 | 0.80 | 0.0075 | 0.0191 | 0.523 |
| RU2 | 0.5 | 180 | 6 | 3 | - | - | - | - |
| SE1 | 2 | 45 | 7 | 2 | - | - | - | - |
| SE2 | 30 | 60 | 6 | 6 | 1.00<sup>H</sup> | 0.0124 | 0.0223 | 0.340 |
| SE3 | 21 | 119 | 9 | 8 | 0.88 | 0.0141 | 0.0263 | 0.340 |
| UA1 | >20,000 | - | 10 | 6 | 0.56 | 0.0083 | 0.0212 | 0.487 |
| UA2 | 2,335 | - | 11 | 7 | 0.60 | 0.0114 | 0.0221 | 0.395 |
| **Total**<br>**(± SE)** | - | **198,800** | **311** | **222** | **0.78** | **0.0115**<br>**(± 0.0426)** | **0.0314**<br>**(± 0.0856)** | **0.248**<br>**(± 0.414)** |

“- “– not calculated/not relevant, H - highest value among populations, L – lowest value among populations

Sampling was conducted under permits from the relevant national authorities and complied with all applicable species protection and conservation regulations.

DNA was extracted using a modified CTAB protocol (Supplementary Data SD1), since *D. rotundifolia* contains high amounts of polyphenols.

### 2.2. Sequencing and data analysis of cpDNA

At first, we tested 18 universal and cross-species plant primer pairs for a set of samples (Supplementary Tables ST2, ST3). Additionally, we designed seven specific primer pairs using the reference plastome of *D. rotundifolia* (Gruzdev et al. 2019; NCBI GenBank: KU168830) and targeting intergenic spacer regions. All specific primers and nine pairs of cross-species successfully amplified and yielded cpDNA sequences. Alignment of obtained sequences revealed no variation.

To detect variable regions in cpDNA and enable design of *D. rotundifolia*-specific primers, a reference-based assembly of the complete plastome was performed. To achieve this, we employed two complementary approaches: (i) long-read sequencing using the Oxford Nanopore platform and (ii) short-read data derived from RNA-seq generated on the Illumina platform.

For the long-read approach, ten samples representing six populations (FI4 (n = 2), DK, DE4, LV8 (n = 3), NO3 (n = 2), SE5) were used. To ensure sequencing of enriched cpDNA, chloroplasts from selected samples were concentrated using the Percoll gradient technique (Supplementary Data SD2). Long-read library preparation was conducted using the Native Barcoding Kit 96 V14 (SQK-NBD114.24; Oxford Nanopore Technologies (ONT), Oxford, UK), following the Ligation Sequencing gDNA protocol: 10 µL of DNA was used for each sample. Sequencing was performed using a FLOW-MIN114 Flow Cell (ONT) and a MinION Mk1C device (ONT; software version 24.11.8). The run duration lasted 100 h. Demultiplexing as well as length filtering (> 100 bp) and quality filtering (> Q10) were done by the MinION device.

In average, we obtained 6,647 long reads per sample (SD ± 2,438). The reads were mapped to the reference plastome using wf-alignment nextflow workflow v1.2.2 (ONT). In average, 1,260 reads were aligned to the reference (SD ± 461) with a mean coverage of 87% (SD ± 4) and a mean sequencing depth of 37 (SD ± 17). Consensus sequences were generated with Medaka v2.0.1 (ONT) and aligned using HAlign v3.0.0_rcl (https://github.com/malabz/HAlign-3).

For the short-read approach, we used RNA-seq data from ten samples representing eight populations (DE1, DE5, DK, FI1, LT1 (n = 2), LV (n = 2), SE3 (n = 2)). RNA extraction, messenger RNA enrichment, and sequencing on a NovaSeq X Plus system (PE150) were performed by Novogene GmbH (Planegg, Germany). In average, 92 M paired reads per sample were obtained (SD ± 8 M). For each sample, a transcriptome was assembled using SPAdes v3.15.4 (Prjibelski et al., 2020) with the parameters --rna and --cov-cutoff off. The resulting transcripts were mapped to the reference plastome using minimap2 with parameters-a-x asm5 (v2.28-r1209; Li 2018) and processed with samtools (view-b-F 4) (v1.21; Danecek *et al*. 2021). Average coverage of 78% (SD ± 0.03, minimum of one transcript per site) of the reference plastome was obtained. Consensus sequences were generated with bcftools (v1.21; Danecek *et al*. 2021) and aligned using HAlign.

Demultiplexed long and short reads are available in GenBank under PRJNA1377308. The resulting alignments were used to design primer pairs targeting the most variable regions to confirm the found SNPs by next-generation sequencing with the Sanger sequencing method: 12 primer pairs were developed based on long-read data and 10 based on short-read data. In total, we sequenced at least three samples for each of 37 primer pairs, covering 10,449 bp (5.42% of 192,912 bp) of the *D. rotundifolia* chloroplast genome. All primer sequences, the tested annealing temperatures and the list of sequenced samples are provided in Supplementary Tables ST2 and ST3. Protocols of PCR (apart from annealing temperatures) and Sanger sequencing were performed according to Kuprina *et al*. (2022; *rbcL–psaI* marker).

### 2.3. Genotyping by ddRADseq

Double digest Restriction-site Associated DNA sequencing (ddRADseq) was performed by LGC Genomics (Berlin, Germany) using PstI-ApeKI restriction enzymes, followed by selection of the fragments 300-500pb long and sequencing on the Illumina NovaSeq 6000 platform (150 bp PE mode, 300 cycles). Samples were arranged quasi-random across four 96-well plates to reduce the batch effect. Six technical replicates were included (the same DNA extract was processed in parallel twice).

Raw reads were demultiplexed and trimmed using bcl2fastq v2.20, allowing two mismatches or Ns in the index barcodes, no mismatches in the inline barcodes, and permitting Ns within the restriction site. Additionally, first six bases of reads were trimmed by trimmomatic v0.39. Read quality was assessed using fastQC v0.12.1 and multiQC v1.32. All reads passed the sequence quality, per base N and adapter content. The multiplexed read sequences for all 317 samples (including six technical replicates) can be found on GenBank under PRJNA1377308. The MultiCQ report is posted on GitHub, https://github.com/kuprinak/Drosera_pogen.

The Single Nucleotide Polymorphisms (SNPs) calling was performed in ipyrad v0.9.105 (Eaton and Overcast 2020) with *de novo* assembly method, datatype “pairddrad”, max cluster depth within samples of 20,000. The optimal assembly parameters were selected by running the analysis with nine different values of “clustering threshold for *de novo* assembly”(ct = 0.83–0.99, with a step of 0.2), two values of “minimum depth for statistical base calling” (md = 6 and 10), and four values of “minimum number samples per locus for output“(ms = 150, 200, 250, 300). In total, 72 obtained datasets were evaluated following the method of McCartney-Melstad *et al*. (2019), and the one obtained with parameters ct = 95, md = 6, and ms = 200 was determined as optimum for the downstream analyses (Supplementary File SF1). After removing 245 loci with more than two alleles, the dataset included 24,852 biallelic SNPs distributed across 7,493 loci with a total aligned length 1,422,473 bp and an average of 19.2% missing data for 317 samples.

### 2.4. Patterns of genetic variation

The ipyrad SNP matrix was used for a Maximum Likelihood phylogenetic reconstruction using IQ-TREE v3.0.1 (Wong *et al*. 2025) with 10,000 ultrafast bootstrap replicates (Hoang *et al*. 2018). The optimal substitution model selected by ModelFinder based on BIC was TN+F+R6. Phylogenetic reconstruction was primarily used to assess the error rate between technical replicates and to identify clonal samples. All six pairs of technical replicates clustered together, with branch lengths between replicates equal to zero in all cases except one (sample UA1-3), where the branch length was 0.001, reflecting 11 mismatches. Accordingly, samples were assigned to the same multilocus genotype (MLG) when the branch length connecting them was equal to or less than 0.001. (Supplementary File SF2). Clonal samples and technical replicates were excluded from subsequent analyses. In addition. All analyses were conducted in R v4.5.0 and Python v3.9.18.

Principal Coordinates Analysis (PCoA) was performed to visualize Euclidean distances among samples with Lingues correction in the R package dartR v2.9.9.5 (Mijangos *et al*. 2022).

To assess clustering patterns with model-based approaches, ADMIXTURE and Discriminant Analysis of Principal Components (DAPC; Jombart et al. 2010) were conducted. We evaluated ancestral cluster numbers (K) from 1 to 20. For each K, we ran ADMIXTURE v1.3.0 (Alexander and Lange 2011) with 10-fold cross-validation (CV) to assess model fit. DAPC was conducted in the R package adegenet v2.1.11 without prior group assignment and with 1,000 principal components retained for cluster detection (n.pca = 1000). To determine the number of genetic clusters, we examined changes in CV for ADMIXTURE and in Bayesian Information Criterion (BIC) values for DAPC across tested K values (Supplementary Figures S1, S2). For both data analyses, we selected K = 2, 3 and 4 for further interpretation.

For subsequent analyses, populations containing fewer than five distinct MLGs were excluded to minimize sampling bias, resulting in a clone-corrected dataset comprising 187 samples from 25 populations (Table 1).

An index of clonal richness C (ranging from 0 to 1) was calculated according to the formula: С = (MLG - 1)/(n - 1) (Dorken and Eckert 2001). Pairwise Fst values and Nei’s distances (D) between populations were calculated in the R package StAMPP v1.6.3 (Pembleton *et al*. 2013). For the Fst estimation, 1000 bootstrap replicates were used. Overall and population-wise estimates of observed heterozygosity (Ho), unbiased expected heterozygosity (uHe) and inbreeding coefficient (Fis) were calculated in R package dartR v2.9.9.5 by the functions “gl.report.heterozygosity”.

The significance of isolation by distance (IBD) was assessed using a Mantel test, based on Pearson’s correlation between a matrix of pairwise genetic p-distances and a matrix of Euclidean geographic distances. The test was performed with 10,000 permutations in the R package vegan v2.7-1 (Oksanen *et al*. 2025).

### 2.5. Isolation by climate

To investigate the influence of climate on genetic variation, we performed a global and a partial redundancy analysis (RDA). Nineteen climatic variables (BIO1–BIO19) were extracted for all sampling locations using bioclimatic layers from WorldClim v2.1 (Fick and Hijmans 2017) at 30 arc-second resolution (∼1 km) and terra v1.8-60 package in R. To reduce collinearity among predictors, pairwise Pearson correlations were calculated between variables, and a subset of weakly correlated variables was retained (r > ±0.75). The final set included BIO1 (annual mean temperature), BIO4 (temperature seasonality), BIO12 (annual precipitation), and BIO15 (precipitation seasonality). These variables were standardized (mean = 0, SD = 1) prior to analysis. No variables exhibited problematic collinearity, as assessed using the vifstep function in the R package usdm v4.5.2 (Naimi *et al*. 2014), with a threshold of 10.

To test whether climatic variation explains patterns of genetic variation, we performed a global redundancy analysis (RDA) using the centered genetic matrix as the response and four selected bioclimatic variables as predictors. To assess the effect of climatic variables while controlling for spatial autocorrelation, we conducted a partial RDA, conditioning on geographic coordinates (longitude and latitude). This analysis allowed us to test for isolation by climate (IBC) independently of IBD. Each RDA was tested with 10,000 permutations using the “rda” function from the R package vegan v2.7-1. To visualize the spatial distribution of genetic variation along the first two RDA axes, site scores for RDA1 were extracted for each individual and plotted on the geographical map using R package sf v1.0-21 (Pebesma and Bivand 2023).

### 2.6. Demographic history

To infer the relative demographic history of *D. rotundifolia* across Europe, we reconstructed changes in effective population size (Ne) through time. Individuals were assigned to one of three genetic clusters based on the highest ancestry proportion inferred by ADMIXTURE. All populations were included, resulting in a total of 222 samples after clone correction (118, 81, and 23 samples for K = 1, 2, and 3, respectively). SNP filtering was performed using VCFtools v0.0.16 (Danecek *et al*. 2011), retaining loci with a maximum missing data proportion of 0.7 and a minimum minor allele count of 2. This filtering step retained 5,049 SNPs. For each cluster, a folded site frequency spectrum (SFS) was generated using the *easySFS* Python script (https://github.com/isaacovercast/easySFS). To account for differences in sample size among clusters, each SFS was projected to 46 chromosomes. The resulting SFSs were used as input for demographic inference in Stairway Plot v2.1.3 (Liu and Fu 2015). In the absence of a species-specific mutation rate for *Drosera*, we assumed a nuclear mutation rate of 1 × 10⁻⁸ mutations per site per generation, consistent with empirical estimates of germline mutation rates in multicellular eukaryotes (Lynch 2010, Wang et al. 2023). Because generation time (average age at reproduction) is difficult to estimate in *D. rotundifolia* due to its perennial and clonal growth habit, we approximated it as one year. Owing to uncertainty in both generation time and mutation rate, our reconstruction focuses on relative changes in effective population size over time rather than precise estimates of absolute timing or Ne values.

## 3. Results

### 3.1. Genetic and clonal diversity

With ddRAD data, we identified 222 unique multilocus genotypes (MLGs) among 311 sampled individuals. Only one population was found to be monoclonal (DE1; n = 4; population diameter = 30 m). Three population pairs shared identical MLGs: LT1 and LT2 (500 m apart), PL5 and PL6 (1,000 m apart), and DE1 and DE2 (6,200 m apart). In some populations each sample represented a unique MLG: DE8, FI2, FI3, FI4, PL1 (Table1, Supplementary Table ST1). The total clonal richness calculated for the populations with more than five samples was 0.78.

Across all populations, the adjusted observed heterozygosity (Ho) was 0.0115 (SD ± 0.0426) and the unbiased expected heterozygosity (uHe) was 0.0314 (SD ± 0.856), resulting in an overall inbreeding coefficient (Fis) of 0.248 (SD ± 0.414) (Table 1). The lowest Ho was observed in population PL2 (0.0071), which also exhibited the highest Fis (0.547). Conversely, the highest Ho (0.0156) was found in FI1, while the lowest Fis occurred in DE8 (0.288). In all populations, observed heterozygosity was lower than the corresponding expected heterozygosity, indicating widespread heterozygote deficiency.

For the cpDNA data, only a single SNP was detected by Sanger sequencing, observed in one sample from population DK at the locus *rps14/psaB*. All the other loci shared the same haplotype for sequenced samples, regardless if SNPs were detected with long ONT reads (1,374 SNPs) or short Illumina reads (482 SNPs).

### 3.2. Genetic differentiation

Principal Coordinates Analysis (PCoA; Figures 2a, 2b) revealed the strongest differentiation of the samples along axis 1, which explained 5.63% of the total genetic variation. Samples from populations DE1-DE5 formed a cluster on the left side of the plot, separated from the rest of the samples, while FI5 positioned at the right-most extreme. Individuals from the same population generally clustered closely together.

**Figure 2.**
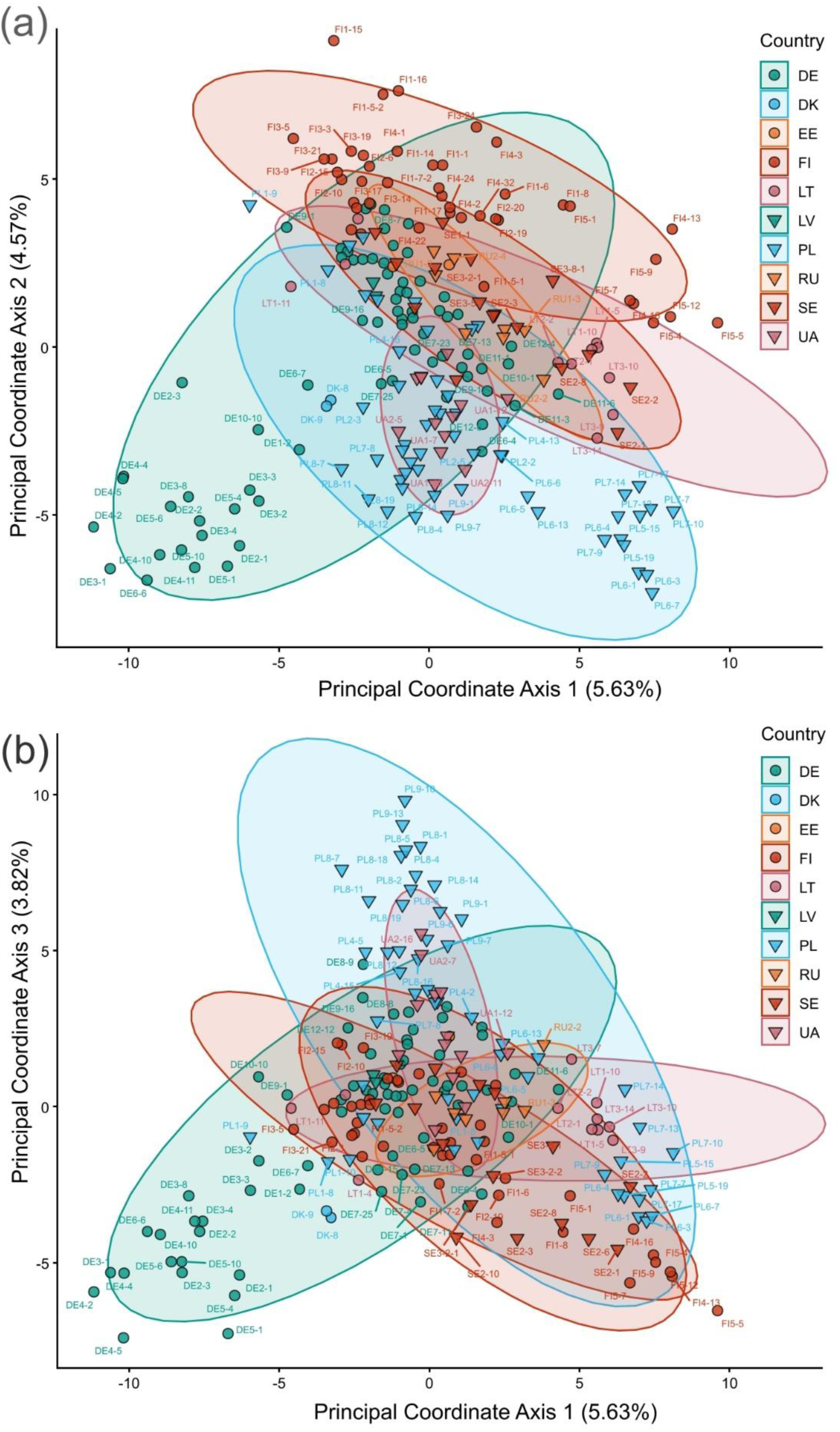
Principal coordinates analysis (PCoA) of Euclidean genetic distances among 222 *Drosera rotundifolia* samples, with 95% ellipses illustrating within-country clustering in ordination space. (a) The top panel shows the top 1 (x-axis) and 2 (y-axis) composite variables, (b) the bottom panel shows the top 1 (x-axis) and 3 (y-axis) composite variables. The dataset comprised 24,852 ddRAD-derived SNPs.

ADMIXTURE and DAPC produced consistent results (Figures 3a-3f). The strongest genetic differentiation, together with the largest changes in CV and BIC values (Supplementary Figures S1, S2), was observed when the dataset was partitioned into two clusters. Cluster 1 comprised the north-western populations from Germany (DE1-DE6) and the population from Denmark (DK). Assuming three clusters (K = 3), ADMIXTURE partitioned samples into three geographically distinct groups: a Western (cluster 1), Northern (cluster 2), and Eastern clusters (cluster 3), with Eastern and Northern clusters being more genetically similar to each other than to Western cluster. Under the four-cluster scenario (K = 4), the Western cluster remained unchanged and the most distinct, whereas the remaining samples were distributed among three weakly differentiated clusters.

**Figure 3.**
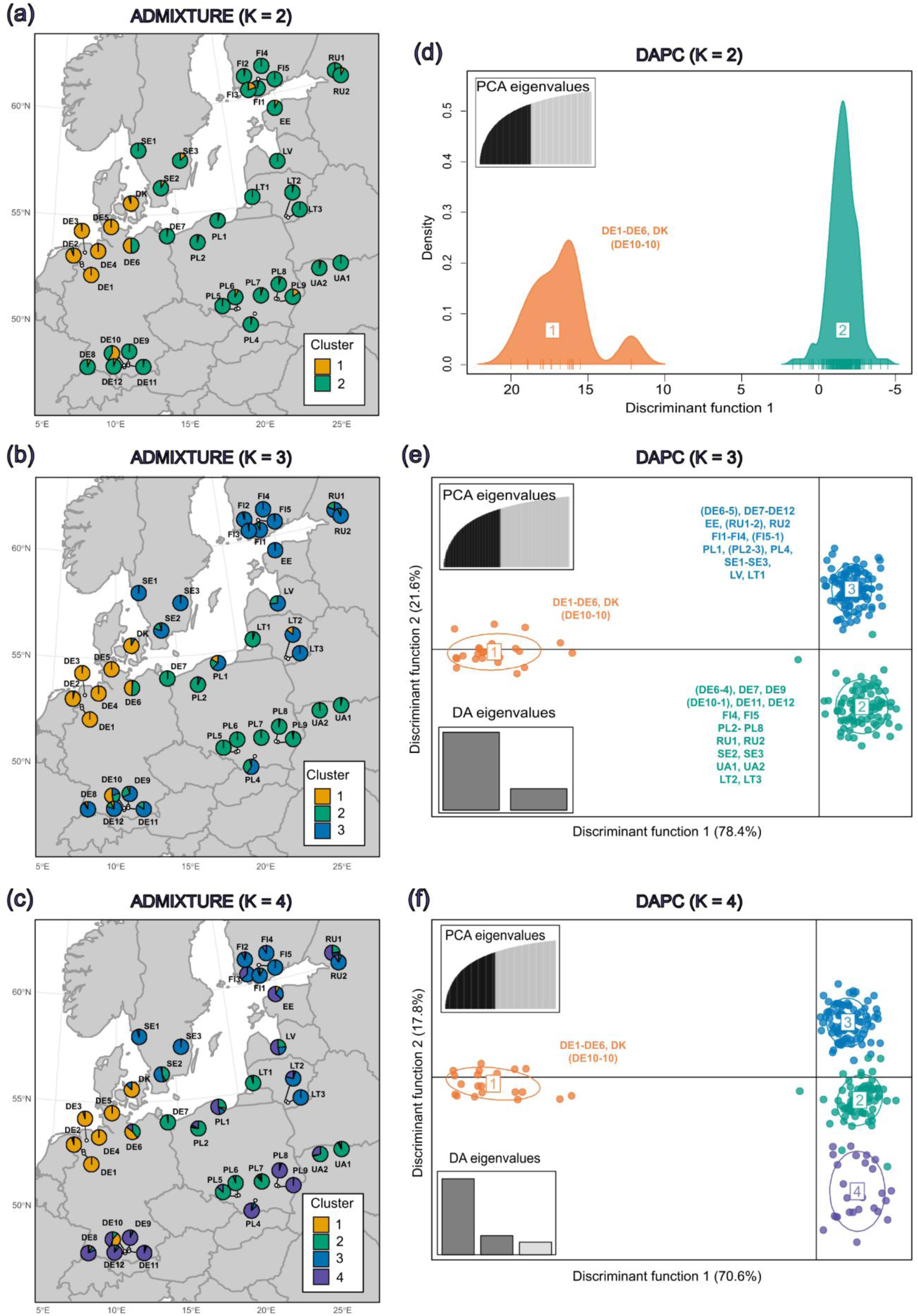
(a, b, c) Geographic distribution of admixture proportions inferred by ADMIXTURE for (a) K = 2, (b) K = 3 and (c) K = 4. (c, d, e) Discriminant Analysis of Principal Components (DAPC) results for (a) K = 2, (b) K = 3 and (c) K = 4. The size of the inertia ellipses encompasses 67% of individuals. Analysed dataset comprised 24,852 ddRAD-derived SNPs from 222 *Drosera rotundifolia* samples from 38 populations.

The Mantel test detected a significant IBD pattern (r = 0.1807, *p* = 0.0001), with genetic distance increasing with geographic distance and explaining 3.3% of genetic variation (Figure 4). We observe significant Fst values (*p* < 0.05) between all compared populations (Figure 5). The highest Fst and Nei’s genetic distances were found between populations FI5 and DE4 (0.44 and 0.026, respectively; Figure 5).

**Figure 4.**
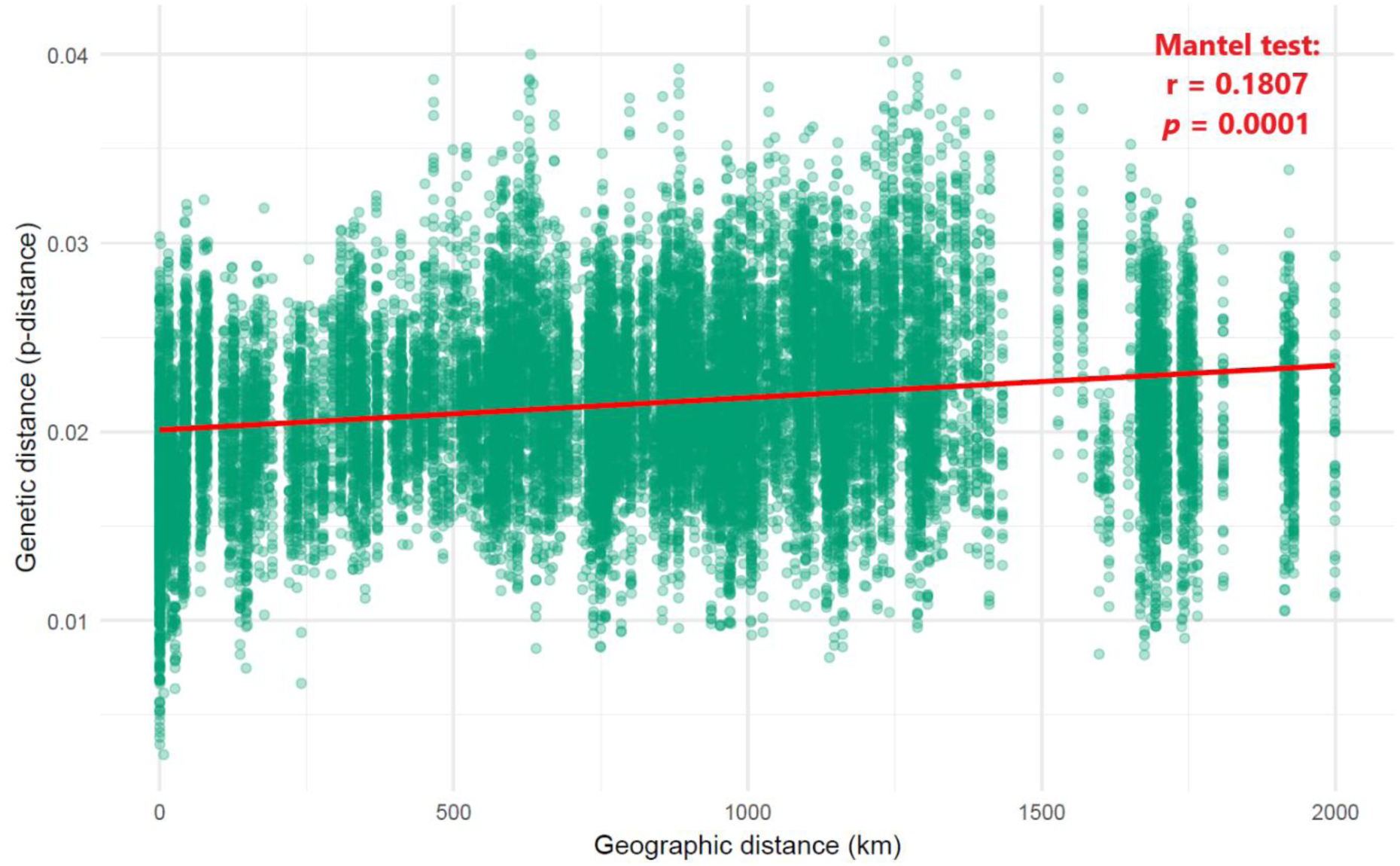
Isolation by distance (IBD) graph between 222 samples of *Drosera rotundifolia*, showing a significant IBD effect (Mantel test, r = 0.1807, *p* = 0.0001). Each point represents a pairwise comparison between two samples; the red line indicates the fitted linear model.

**Figure 5.**
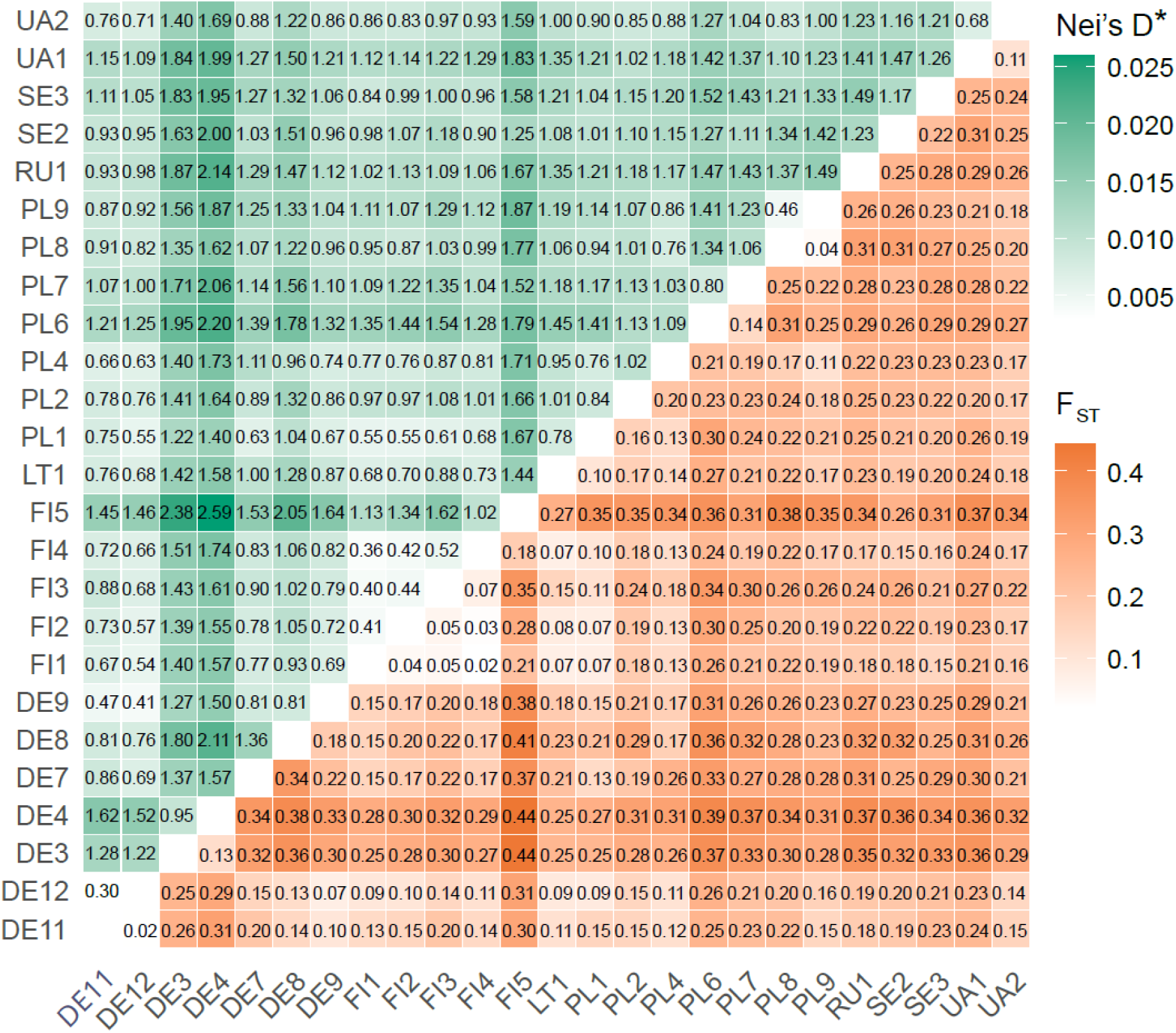
Heatmap of pairwise population comparisons showing Fst values (lower triangle) and Nei’s genetic distances (D, upper triangle) between 25 populations of *Drosera rotundifolia* after clone correction. *Nei’s D values on the heatmap were multiplied by 100 for visualisation purposes. All *p*-values for Fst < 0.05.

### 3.3. Isolation by climate

The global RDA indicated that tested climatic variables explained a significant portion of genetic variation across populations of *D. rotundifolia*. The overall model was significant (permutation test, p < 0.0001 for each predictor), with an adjusted R² of 0.061, indicating that 6.1% of the variation in the genetic matrix was explained by the four selected bioclimatic variables.

After controlling for special effects, the partial model remained significant (permutation test, *p* < 0.0001), with an adjusted R² of 0.029. This indicates that climatic variables explained 2.9% of the genetic variation, regardless of special structure. All four predictors remained significant in the partial RDA (BIO1 - annual mean temperature): F = 3.73, *p* < 0.0001; BIO4 – temperature seasonality: F = 1.82, *p* = 0.0004; BIO12 - annual precipitation: F = 2.56, *p* < 0.0001; BIO15 – precipitation seasonality: F = 2.64, *p* < 0.0001). In total, 1,022 SNPs from 632 loci were found to be correlated with the climate variables.

First two axes of RDA explained 3.2% of the total genetic variation after conditioning on geography (Figure 6a) with annual mean temperature and annual precipitations having the strongest influence. The spatial distribution of RDA1 values revealed three geographically coherent clusters that correspond closely to the ADMIXTURE results at K = 3 (Figure 6b). Populations from FI, SE, and PL1 showed the lowest RDA1 scores, which are associated with cooler annual temperatures, higher annual precipitation, and greater temperature and precipitation seasonality. In contrast, samples from the south-eastern population exhibited the highest RDA1 scores, indicating association with warmer and less seasonal climatic conditions.

**Figure 6.**
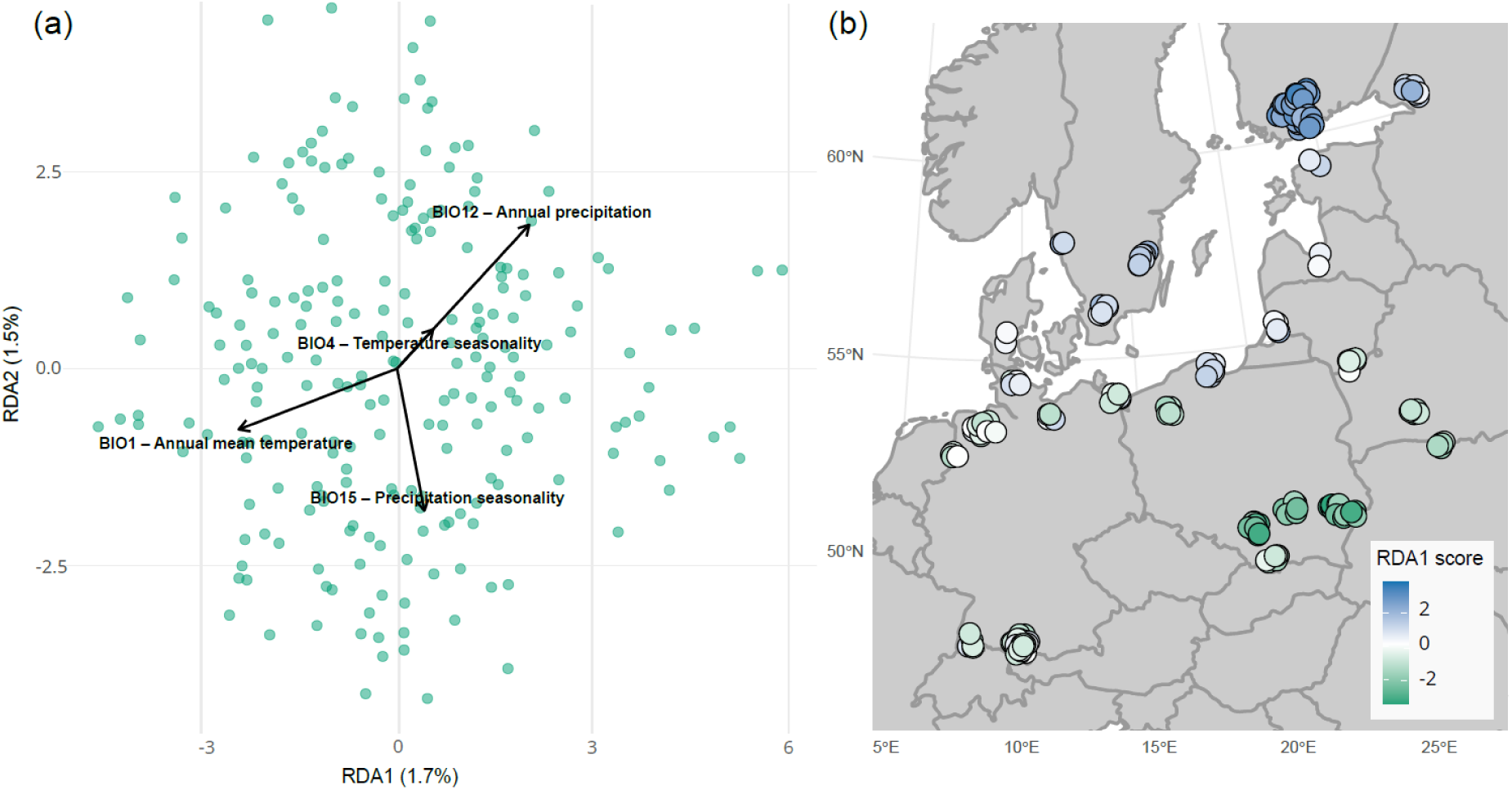
(a) Partial Redundancy Analysis (RDA) biplot showing the first two constrained axes (RDA1 and RDA2) and their relationships with four climatic variables (BIO1, BIO4, BIO12, BIO15) after conditioning on geographic coordinates for 222 samples of *Drosera rotundifolia* from 38 populations. (b) Spatial distribution of individual scores along global RDA1 across Europe. Sample points are jittered for visualisation purpose.

### 3.4. Demographic history

The Stairway Plot analysis (Figures 7a, 7b, 7c) revealed historical expansions in effective population size (Ne) across all three genetic clusters. Cluster 1 (West) experienced the most recent and pronounced expansion event, occurring ca. 500–900 generations ago. Following a severe decline to near-zero levels, Ne increased to ca. 300,000 individuals. In contrast, clusters 2 (East) and 3 (North) exhibited earlier demographic expansions beginning ca. 10,000 generations ago from an ancestral Ne of ca. 8,000 individuals. Cluster 2 experienced two successive expansion phases, each resulting in a fivefold increase in Ne: the first occurred ca. 12,000-6,000 generations ago, whereas the second took place ca. 2,500-3,500 generations ago. Cluster 3 underwent a single expansion event ca. 6,000-10,000 generations ago, resulting in a ca. tenfold increase in Ne.

**Figure 7.**
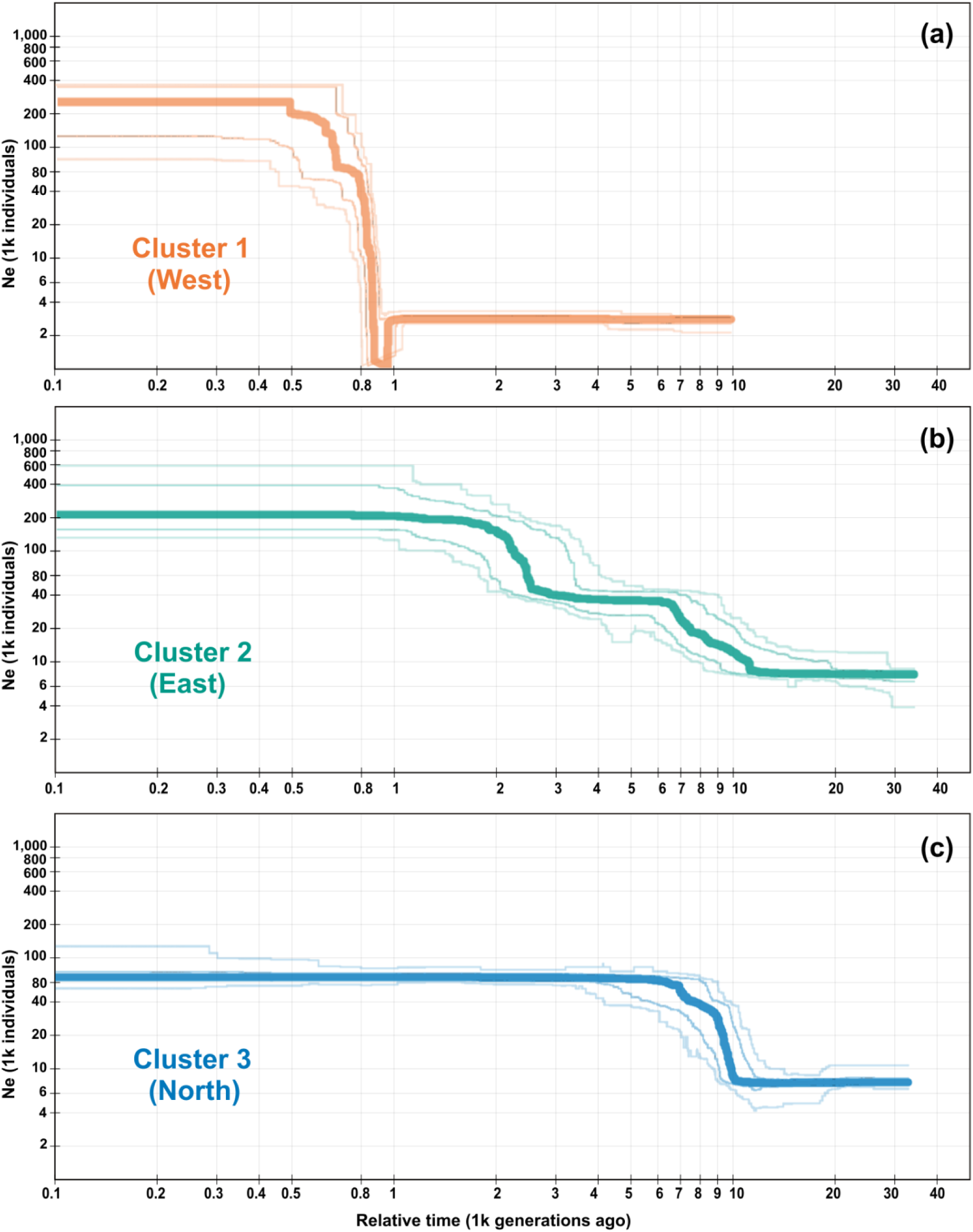
Historical changes in effective population size (Ne) of three European genetic clusters of *Drosera rotundifolia* inferred using Stairway Plot 2. Individuals were assigned to clusters based on ADMIXTURE analysis with K = 3. Panels show demographic histories for (a) cluster 1 (West), (b) cluster 2 (North), and (c) cluster 3 (East). Middle line in each panel - median of 200 inferences based on subsampling. Light lines: 75% (inner) and 95% (outer) confidence intervals of the inference.

The contemporary effective population size of cluster 3 remained smaller (Ne ≈ 80 × 10³) than that of clusters 1 and 2. No pronounced changes in Ne were detected in any cluster during the last 500 generations.

## 4. Discussion

### 4.1. Clonal variation

Vegetative reproduction in *D. rotundifolia* appears to be more widespread than expected for a species with the dominant reproduction mode as sexual and which does not form long rhizomes or stolons. New buds develop on the adaxial surface of the lamina, on the petiole, in the axils of the leaves and on the flower stalks, as well as on the root suckers (Baranyai and Joosten 2016). Approximately 70% of all sampled individuals were unique MLGs, resulting in an overall clonal richness of 0.78. In one small population (DE1; ∼30 m in diameter), only a single MLG was detected. Taking into account the error rate of our genotyping method, we considered two samples to represent distinct MLGs when they differed by more than 11 mismatches among 24,852 analysed SNPs.

Long-distance dispersal of clones in *D. rotundifola* appears uncommon but is possible. The greatest distance between genetically identical samples within a single population was 490 m, observed in population FI1 (MLG 78; Appendix Table 1). Three pairs of populations shared identical MLGs: LT2 and LT3 (500 m apart), PL5 and PL6 (1,000 m apart), and DE1 and DE2 (6,200 m apart). While it is possible that ramets were transferred by birds or humans between populations, alternative explanations include fragmentation of the bog in the past (e.g., due to human settlements in PL or DE or forest succession in LT).

How fast a genotype of *D. rotundifolia* can disperse clonally? In a two-year study of a subarctic bog in Sweden, Svensson (1995) estimated an average of 1.57 ± 0.8 ramets per genet (maximum 4; SD, n = 93) of *D. rotundifolia*. This corresponds to an approximate production rate of 0.3 ramets per genet per year. To spread across a bog with a diameter of only 30 m, such as DE1, a single genotype would require over 1,000 years (assuming each clonal propagation event results in dispersal of ca. 5 cm, expansion from the population centre, and no artificial transfer of clones). It is hard to assume that no mutations would occur over such a timescale. There is no evidence for particularly low mutation or recombination rates in *Drosera*, though it would be interesting to investigate.

### 4.1. Genetic variation: cpDNA

We detected exceptionally low variation in the chloroplast DNA (cpDNA) of *Drosera rotundifolia*. Sanger sequencing of 10,449 bp of the plastome, while targeting potentially the most variable regions, identified only a single SNP in one locus.

Amplification of cpDNA using universal and cross-species primers was largely unsuccessful, whereas all *D. rotundifolia*-specific primers yielded PCR products. This observation points towards the extensive plastome rearrangements and gene losses which are characteristic of carnivorous plants and Droseraceae (Gruzdev *et al*. 2019; Nevill *et al*. 2019; Fu *et al*. 2023). In *D. rotundifolia*, these include the loss of *trnA-UGC*, *trnV-UAC*, *ycf1* and all 11 *ndh* genes. In contrast to the generally compact plastomes of many carnivorous plants, *D. rotundifolia* possesses a relatively large plastome (∼192,912 bp), mostly due to expansion of the inverted repeat (IR) region. Its plastome is highly rearranged and repeat-rich, comprising 23.13% repetitive sequence and containing an exceptionally high number of repeats larger than 20 bp (1,089).

To date, no published data are available on interspecific variation in *Drosera* cpDNA. The only relevant study is a phylogenetic analysis by Rivadavia *et al*. (2003), which found that *rbcL* sequences of *D. rotundifolia* and *D. anglica* differed by a single nucleotide, suggesting that *D. rotundifolia* can be the maternal parent of *D. anglica*.

RNA-seq data have previously enabled complete chloroplast genome assembly in three *Erysimum* (Brassicaceae) species (Osuna-Mascaró *et al*. 2018) and appeared promising for *D. rotundifolia* given the low level of RNA editing reported for *D. rotundifolia* (only six confirmed editing sites compared to ca. 30–40 typically found in angiosperms; Gruzdev et al. 2019). Rather than mapping reads directly to the reference plastome, we first assembled the transcriptome, which in our experience resulted in higher overall coverage and lower misassemble rates. However, RNA-seq data did not yield a complete plastome assembly either way. Average coverage reached 78% of the reference plastome, which can be compared to the 78% plastome exon coverage reported by Gruzdev *et al*. (2019).

Although our next-generation sequencing approaches detected variation, Sanger sequencing confirmed only a single SNP. At this stage, we cannot conclude that cpDNA variation in *D. rotundifolia* is entirely absent. Both sequencing approaches have inherent limitations: Oxford Nanopore long reads are associated with relatively higher error rates (Bejaoui *et al*. 2025), while short-read Illumina data can be difficult to map accurately in highly repetitive regions, such as those abundant in the *D. rotundifolia* plastome (Fu *et al*. 2023). Future studies employing higher-accuracy long-read technologies, such as the PacBio platform, may provide a more reliable assessment of cpDNA variation in *D. rotundifolia*.

### 4.2. Genetic variation: ddRADseq

All studied populations of *D. rotundifolia* exhibited low genetic variation and a pronounced deficit of heterozygosity (overall uHe = 0.0314, Ho = 0.0015, Fis = 0.248). The lowest observed heterozygosity (Hₒ = 0.0071) was recorded in population PL2, which also exhibited the highest inbreeding coefficient (Fis = 0.547). This population occupies a 7.2 ha bog within a forest. Such small, isolated populations are particularly vulnerable, being prone to increased homozygosity and inbreeding. This can lead to inbreeding depression (Charlesworth and Willis 2009) and ultimately compromise population viability (Frankham 2005).

These estimates show markedly lower genetic diversity level than those reported in previous studies of *D. rotundifolia*: Eschenbrenner *et al*. (2019) documented a minimum He of 0.109 and a maximum of 0.247 using ISSR markers, while Chung *et al*. (2013) reported a total He of 0.062 based on allozymes. While overall higher Fis values were found for four populations in North Korea, (0.563), even higher Fis values were found for some of European populations, for example, in PL2 (0.547), DE4 (0.493) and DE3 (0.455) (Table 1).

As no ddRAD data are currently available for any *Drosera* species, we compare our results with those from another European wetland plant, *Viola uliginosa* Besser. This perennial species shares key reproductive traits with *Drosera*, including the ability to reproduce clonally and to produce both chasmogamous and cleistogamous flowers (Małobęcki *et al*. 2016; Paul *et al*. 2016). *Viola uliginosa* inhabits rare eutrophic swamp forests and flooded meadows, habitats characterized by a high degree of population patchiness. In most of the European countries this plant is legally protected. Lee *at al.* (2020) reported inbreeding coefficients (F_SNP_) from Belarus, Poland, Estonia, and Slovenia broadly comparable to those observed in our study (F_SNP_ = 0.407–0.593 vs. FIS = 0.288–0.547), although populations of *V. uliginosa* from Finland exhibited substantially higher values (F_SNP_ = 0.872–0.945) than any studied population of *D. rotundifolia*.

Caution is needed when comparing these studies directly, as the present study employed ddRADseq data with bias correction following Nei (1978), whereas Eschenbrenner *et al*. (2019) and Chung *et al*. (2013) relied on dominant markers corrected by Lynch and Milligan (1994). Even when using similar marker types, differences in loci sets and SNP-calling parameters can yield non-comparable estimates. Consequently, measures of genetic diversity and inbreeding are most reliably interpreted within, rather than across, studies.

### 4.3. Genetic differentiation

Observed patterns of genetic differentiation corresponded to geographic distance, but not in a strictly linear manner. Although the Mantel test indicated a significant isolation by distance (r = 0.18, *p* = 0.0001), and pairwise Fst values revealed significant genetic differentiation among all studied populations (Figure 5), the highest Fst value was observed between populations FI5 and DE4 (ca. 600 km apart), rather than between the most geographically distant populations (e.g., RU1 and DE8, ca. 2,000 km apart).

Clustering analyses based on DAPC and ADMIXTURE consistently separated samples into two major, geographically distant clusters: Western and Eastern. Within the Eastern cluster, differentiation appeared weaker, with subdivision toward Northern and Eastern clusters. The Western cluster (cluster 1) comprised five German populations (DE1–DE6) and a single studied population from Denmark (DK). Only DE6, the easternmost population from the cluster, showed admixture, with individuals assigned to both Western and Eastern clusters; the remaining populations were almost entirely assigned to the Western cluster. These populations vary in size, ranging from 1.5 ha (DE3) to 708 ha (DE2), but are all located within the same geographic region, northwestern Germany and Denmark.

We did not estimate Fst values and diversity indices for all sampled populations due to limited sample sizes and the potential sampling bias. However, sufficient sampling was available for two populations from the Western cluster (DE3 and DE4). These populations exhibited overall the highest pairwise Fst values relative to other populations. The lowest Fst was observed between these populations and PL1 (0.25 and 0.27, for DE3 and DE4, respectively), whereas the highest values were found against FI5 (0.44, for both DE3 and DE4). While the Western cluster was the most divergent group of populations, the population FI5 from Finland had the highest Fst values overall, being the most genetically divergent population in our study.

The pronounced genetic divergence of the Western cluster could potentially reflect processes such as cryptic speciation or hybridisation. For instance, *Drosera anglica* Huds. was recently shown to have originated from allopolyploidisation of a hybrid between *D. rotundifolia* and *D. linearis* and it is morphologically distinct from its parental taxa (Mohn and Yang 2026). In contrast, our observation of leaf morphology did not reveal clear differentiation among clusters (Supplementary Figure S4). However, a more detailed examination of morphological, cytological and molecular data could provide further insights.

### 4.4. Isolation by climate

The genetic structure of *D. rotundifolia* was associated with climatic gradients, with 6.1% of the genetic variation explained primarily by mean temperature and precipitation. After accounting for geographic structure, climatic variables still explained a small but significant proportion of genome-wide variation (adjusted R² = 0.029, *p* < 0.0001). This pattern is consistent with polygenic responses to climatic gradients, as 1,022 SNPs from 632 loci were significantly associated with environmental variables. However, the absence of an annotated or chromosome-level reference genome for *D. rotundifolia* is limiting robust genotype–environment association analyses.

Analysis of the geographic distribution of individual RDA1 scores across Europe demonstrated a clear association with three principal biogeographic regions: Atlantic, Boreal, and Continental (Figure 8a). Notably, this pattern is consistent with the clustering inferred by ADMIXTURE and DAPC, which also delineate three main genetic groups (Figure 8b). This concordance suggests that the observed clusters may, at least in part giving only 2.9% of genetic variation explained by climate, reflect adaptation to distinct climatic conditions.

**Figure 8.**
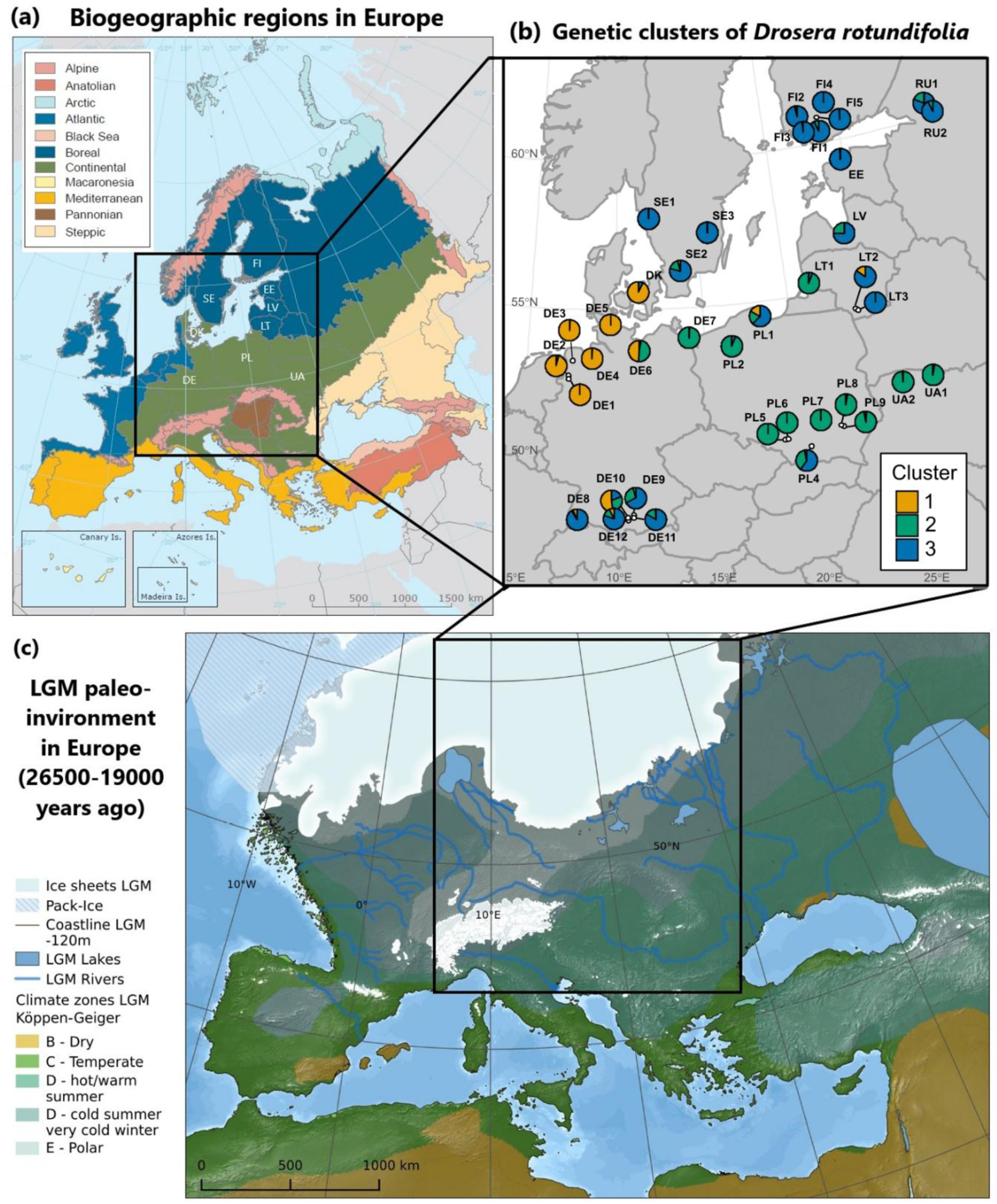
(a) Biogeographical regions of Europe (modified from EEA 2018); (b) Geographic distribution of admixture proportions inferred by ADMIXTURE for K = 3 for 38 populations of *Drosera rotundifolia*; (c) Map of Europe during Last Glaciation Maximum (LGM; modified from Becker *et al*. 2015)

### 4.5 Demographic history

*Drosera rotundifolia* is native to Europe and is hypothesized to have dispersed from South and North America and subsequently via Asia (Rivadavia *et al*. 2003). In Northern Europe, peatlands began to develop ca. 10,000 years ago following deglaciation after the last glacial period (Ruppel *et al*. 2018). Consequently, the formation of the three observed genetic clusters in *D. rotundifolia* is likely influenced by relatively recent historical and demographic processes, like bottlenecks associated with long-term isolation and founder effects due to postglacial recolonization.

Inferring demographic and historical processes from genetic data remains challenging, as bottlenecks and long-term isolation within refugia can produce similar genetic signatures (Stewart *et al*. 2010). Moreover, our time scale should not be interpreted in absolute terms, as no mutation rate or generation time estimates are currently available for *D. rotundifolia*. This perennial species can live for up to five years and can flower starting with their first growing season (Crowder *et al*. 1990). However, extensive clonal growth, mixed mating through self-and cross-pollination, and the persistence of seed banks likely result in complex life-history dynamics.

Nevertheless, it is notable that the Northern genetic cluster of *D. rotundifolia* largely coincides with the area formerly covered by the Fennoscandian ice sheet during the Last Glacial Maximum (LGM) (Figure 8c). In contrast, the Western and Eastern clusters are distributed to the west and east of the periglacial steppe area connecting the Fennoscandian and Alpine ice sheets (Meschede and Warr 2019). Additionally, populations in southern Germany exhibit admixture among all three genetic clusters, suggesting secondary contact rather than persistence in local microrefugia. This pattern is consistent with findings from other peatland species, such as *Rubus chamaemorus* L., where southern populations in Poland and Germany were interpreted as the result of long-distance colonization rather than glacial relicts (Ehrich *et al*. 2008).

The expansion events detected in the Northern and Eastern clusters, both occurring ca. 6,000–10,000 generations ago, were likely associated with the colonization of newly formed peatlands following deglaciation. The second, more recent expansion event identified only in the Eastern cluster ca. 2,500–3,500 generations ago may reflect a second recolonization wave. One possible explanation for the pronounced Ne expansion detected in the Western cluster is a founder event associated with relatively recent colonization ca. 500–900 generations ago, potentially originating from western or southwestern refugial populations, such as those from the Iberian Peninsula (Castro *et al*. 2015; Souto *et al*. 2019). Alternatively, long-distance dispersal, including possible human-mediated introduction, cannot be excluded. Further sampling of *D. rotundifolia* populations from Western and Southwestern Europe, as well as other parts of its range, will be necessary to identify potential source populations. Integrating palaeoecological evidence, such as pollen and macrofossil records, may provide additional insights into the postglacial history of this species.

### 4.6 Conservation strategy

*Drosera rotundifolia* is classified as Least Concern on the European IUCN Red List (Maiz-Tome 2016). However, it is considered vulnerable and legally protected in many European countries and regions since its primary habitat, *Sphagnum*-dominated peatlands, is declining across Europe (Baranyai and Joosten 2016; Germany 2005; Topić and Stančić 2006).

Recent modelling of *Drosera* species in South America suggests that future climate scenarios may pose substantial challenges, with projected declines in habitat suitability under changing temperature and precipitation regimes. Approximately 71.79% of *Drosera* species are expected to experience habitat contraction by 2050–2070 (Olivares-Pinto *et al*. 2025). Given similar climatic trends and ongoing peatland loss in Europe, comparable or higher risks are anticipated for *D. rotundifolia* across its European range.

In light of these findings, adaptive conservation strategies are needed to mitigate the impacts of habitat loss. Consistent with previous recommendations (Eschenbrenner *et al*. 2019; Baranyai and Joosten 2016; Chung *et al*. 2013), such strategies should prioritize protection of existing habitats and preservation of genetic diversity to enhance long-term species resilience. Overall, conservation efforts should explicitly account for the genetic structure of *Drosera rotundifolia*. In particular, seed collection and *ex situ* cultivation should encompass all three major genetic clusters to capture existing evolutionary diversity. Given the potential for local adaptation, assisted migration across distinct climatic regions should be avoided.

Populations with high clonal diversity (e.g., FI2–FI4, PL1, SE2, DE8) represent important genetic resources and should therefore be prioritized for conservation. Conversely, populations exhibiting high inbreeding coefficients and low clonal diversity (e.g., DE3, DE4, DE7, PL2, PL4, RU1) require targeted management interventions, such as assisted migration via regional seed or plant transfer. Additionally, populations DE1–DE6 and DK, which constitute a distinct Central European genetic lineage, represent a particularly important genetic resource and warrant specific conservation attention.

## Conclusions

This study examines the population genomics of *D. rotundifolia* across Europe and reveals three major genetic clusters that broadly correspond to biogeographic regions. Our results indicate that both historical and environmental factors shape the species’ genetic structure. While geographic distance partly explains patterns of genetic differentiation, climatic variables also contribute significantly, consistent with a polygenic response to environmental gradients. The spatial distribution of clusters further suggests that postglacial recolonization and long-term isolation have played a key role in shaping current patterns of diversity.

In contrast to nuclear genomic data, chloroplast DNA exhibited extremely low variation, highlighting its limited utility for phylogeographic inference in this species. In addition, clonal reproduction appears more widespread than expected for a predominantly sexual plant, as reflected by moderate overall clonal richness.

From a conservation perspective, it is essential to preserve genetic diversity across all three major clusters, with particular attention to the Western cluster, while ensuring that management strategies account for potential local adaptation.

## Data Accessibility

Metadata accompanying the data quality and data analysis is posted on GitHub, https://github.com/kuprinak/Drosera_pogen. The demultiplexed fastq data for long and short reads are archived in the NCBI SRA (BioProject ID: PRJNA1377308)

## Acknowledgements

The authors thank all relevant authorities for granting permits for the collection of *Drosera rotundifolia* samples. We are grateful to Rivne State Reserve and Cheremske Nature Reserve, as well as Olga Denyshchyk from the Michael Succow Foundation, for providing plant samples from Ukraine. We thank Gerald Jurasinski, Johann Fermum, Andreas Haberl, Regina Neudert and John Couwenberg from the Greifswald Mire Centre for the help with the fieldwork. Sampling in Finland was supported by Niko Silvan, Bertalan Galambosi and Leila Korpela from the Natural Resources Institute Finland (Luke), as well as by Kari Minkkinen from the University of Helsinki. We also thank and Jūratė Sendžikaitė and Nerijus Zableckis from the Nature Research Centre and the Foundation for Peatland Restoration and Conservation (Vilnius) for their organizational help. We are grateful to Gustaf Granath from Uppsala University, Dominik Zak from Aarhus University, Ilze Ozola from the Latvian National Peatland Society for their support with field sampling in Sweden, Denmark and Latvia. We are also thankful to the people from biological station Osterholz and Förderverein Grambower Moor for guidance in the field.

We thank Pauline Hampe for the meticulous extraction of DNA from the tiny sundew leaves and Oleg Shchepin for his assistance with the ipyrad pipeline.

## Author contribution

KK: Conceptualization, Investigation, Data curation, Data analysis, Visualization, Writing - original draft. MS: Investigation, Data analysis, Visualization, Writing - original draft. HK: Investigation, Writing - original draft. MZ: Investigation, Data curation, Writing - original draft. MS: Conceptualization, Writing - review and editing, Funding acquisition. MB: Conceptualization, Writing - review and editing.

## Benefit-Sharing Statement

The Nagoya Protocol does not apply to this study since no benefits were generated. All applicable species protection and nature conservation regulations were complied with. Regulatory approval for the sampling was obtained from the relevant authorities in the respective countries. To date, *Drosera rotundifolia* is not listed in CITES.

## Funding

This research was funded by the Federal Ministry of Food and Agriculture (BMEL), Agency for Renewable Resources (FNR), Funding code: 2221MT012X. Open Access funding enabled and organized by Projekt DEAL.

## Conflict of Interest

Authors have no conflict of interest to declare.

## Notes

https://github.com/kuprinak/Drosera_pogen

### Competing Interest Statement

The authors have declared no competing interest.

### Summary of Updates

Title updated; Abstract updated; Figure 1 revised; Acknowledgements updated.

