## Supplementary material for "“Population genomics reveals divergent lineages across Europe in the vulnerable mire plant *Drosera rotundifolia”*": SD1

**Supplementary Data SD1.** Protocol of DNA extraction

Dried leaf material (5-10 mg) was ground in a 2-mL tube with a 5-mm steel ball at 3.0 rps for 2 min. Lysis was performed in 600 μL CTAB buffer (pH 8.0) containing 2% CTAB, 100 mM Tris-HCl (pH 8.0), 20 mM EDTA, 1.4 M NaCl, 1% PVP, and 1% β-mercaptoethanol, incubated for 1 h at 65 °C with shaking at 300 rpm. After lysis, samples were centrifuged for 10 min at 6000 rpm, and the supernatant was purified twice by adding 600 μL chloroform:isoamyl alcohol (24:1, v/v), shaking at 300 rpm for 2 min, centrifugation for 10 min at 6000 rpm, and transfer of the aqueous phase to a new tube. The recovered supernatant was mixed with 400 μL isopropanol, shaked at 300 rpm for 2 min, and incubated for 1 h or overnight at –20 °C. Samples were then centrifuged for 10 min at 6000 rpm, and the supernatant was discarded. The DNA pellet was washed twice with 500 μL of 75% ethanol, each time centrifuged for 10 min at 6000 rpm and the liquid removed. Finally, the pellet was air-dried for 1 h at room temperature and resuspended in 100 μL TE buffer.
