## Supplementary material for "“Population genomics reveals divergent lineages across Europe in the vulnerable mire plant *Drosera rotundifolia”*": SD2

**Supplementary Data SD2.** Percoll gradient technique for chloroplast concentration

To ensure sequencing of enriched cpDNA, chloroplasts from selected samples were concentrated using Percoll gradient technique. 50 mL of chloroplast isolation buffer (CIB, pH = 6.1, HCl) with BSA (0.1% w/v) was prepared by dissolving 3.5 g sorbitol, 0.89 g Tris (pH = 7.8), 0.05 g $MgCl_2$*$6H_20$, 0.03 g NaCl, 0.03 g EDTA and 0.05 g BSA in $ddH_20$. Two fresh leaves were chopped in 1.3 mL CIB on ice and transferred to a 2 mL centrifuge tube through a 20 µm filter. Subsequently, samples were centrifuged for 2.5 min at 1000 rcf, while a 40% Percoll solution was prepared in a set of 1.5 mL Eppendorf LoBind® Tubes. The supernatant of the centrifuged samples was removed, leaving a green pellet, which was resuspended in 0.5 mL CIB by carefully pipetting up and down. The 40% Percoll solution (0.4 mL Percoll with 0.6 mL CIB) was then overlaid with the suspension and centrifuged at 1500 rcf for 4 min. The suspension of the centrifuged samples was removed, while the pellet was resuspended in 300 µL CTAB buffer for subsequent DNA extraction. DNA extraction was conducted as in 2.2. Fluorometric measurements (DeNovix, Wilmington, Delaware, USA) indicated a DNA concentration below 10 ng/µl for each sample.
