## Supplementary material for "“Population genomics reveals divergent lineages across Europe in the vulnerable mire plant *Drosera rotundifolia”*": S1

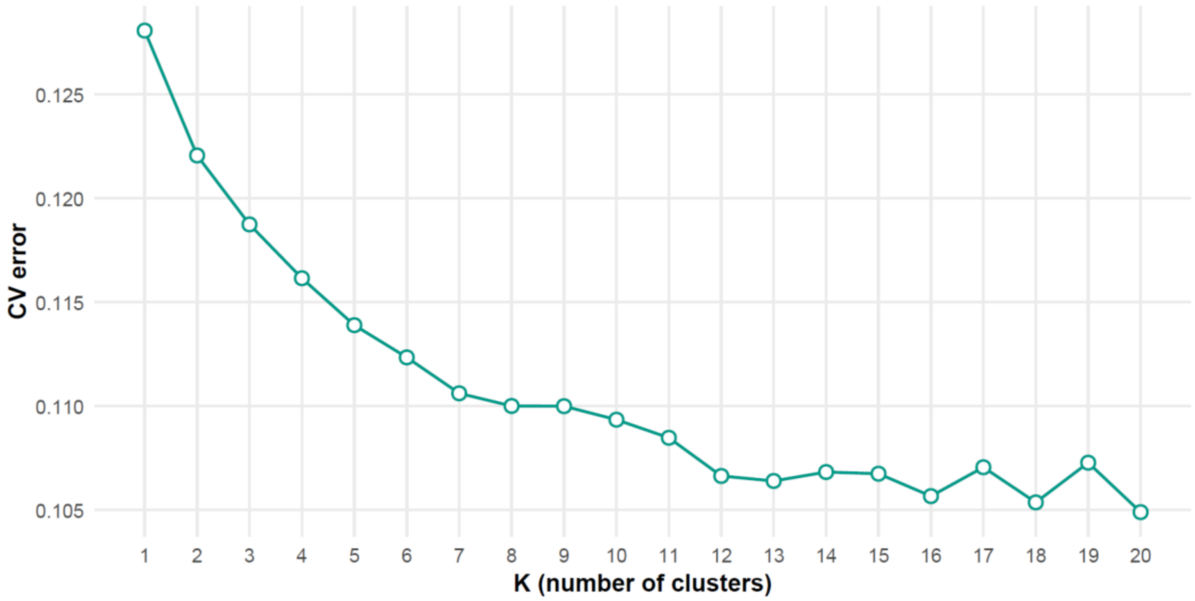


**Supplementary Figure S1.** Cross-validation (CV) error values calculated with 10-fold for 20 clusters (K) in ADMIXTURE. Dataset included 24,852 ddRAD-derived SNPs from 311 samples of *Drosera rotundifolia*
