## Supplementary material for "“Population genomics reveals divergent lineages across Europe in the vulnerable mire plant *Drosera rotundifolia”*": S2

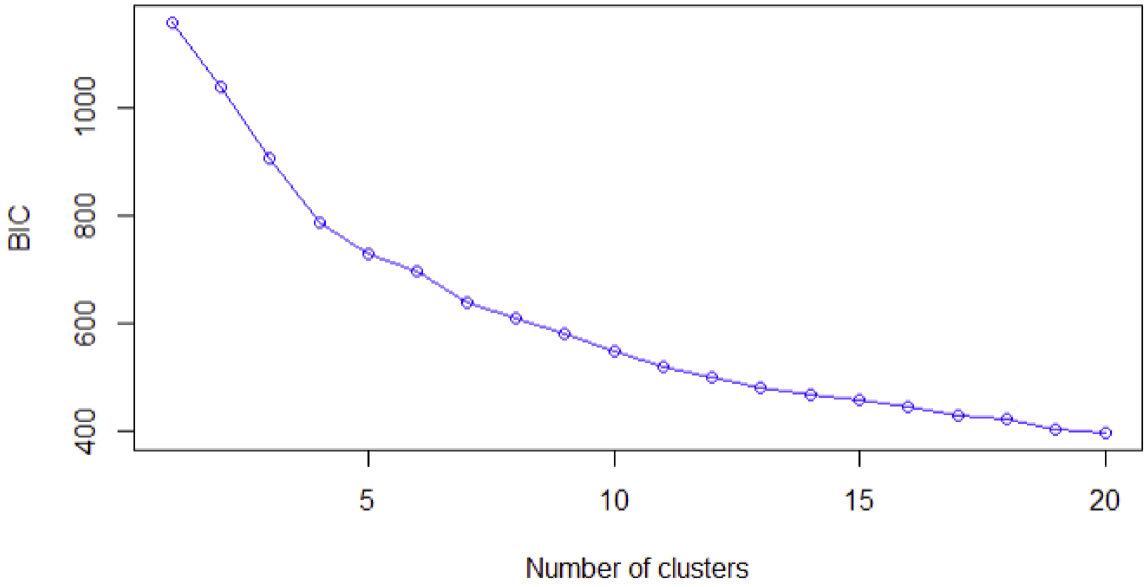


**Supplementary Figure S2.** Bayesian Information Criterion (BIC) values across tested numbers of genetic clusters, calculated using Discriminant Analysis of Principal Components (DAPC). Dataset included 24,852 ddRAD-derived SNPs from 222 samples of *Drosera rotundifolia*
