## Supplementary material for "“Population genomics reveals divergent lineages across Europe in the vulnerable mire plant *Drosera rotundifolia”*": S3

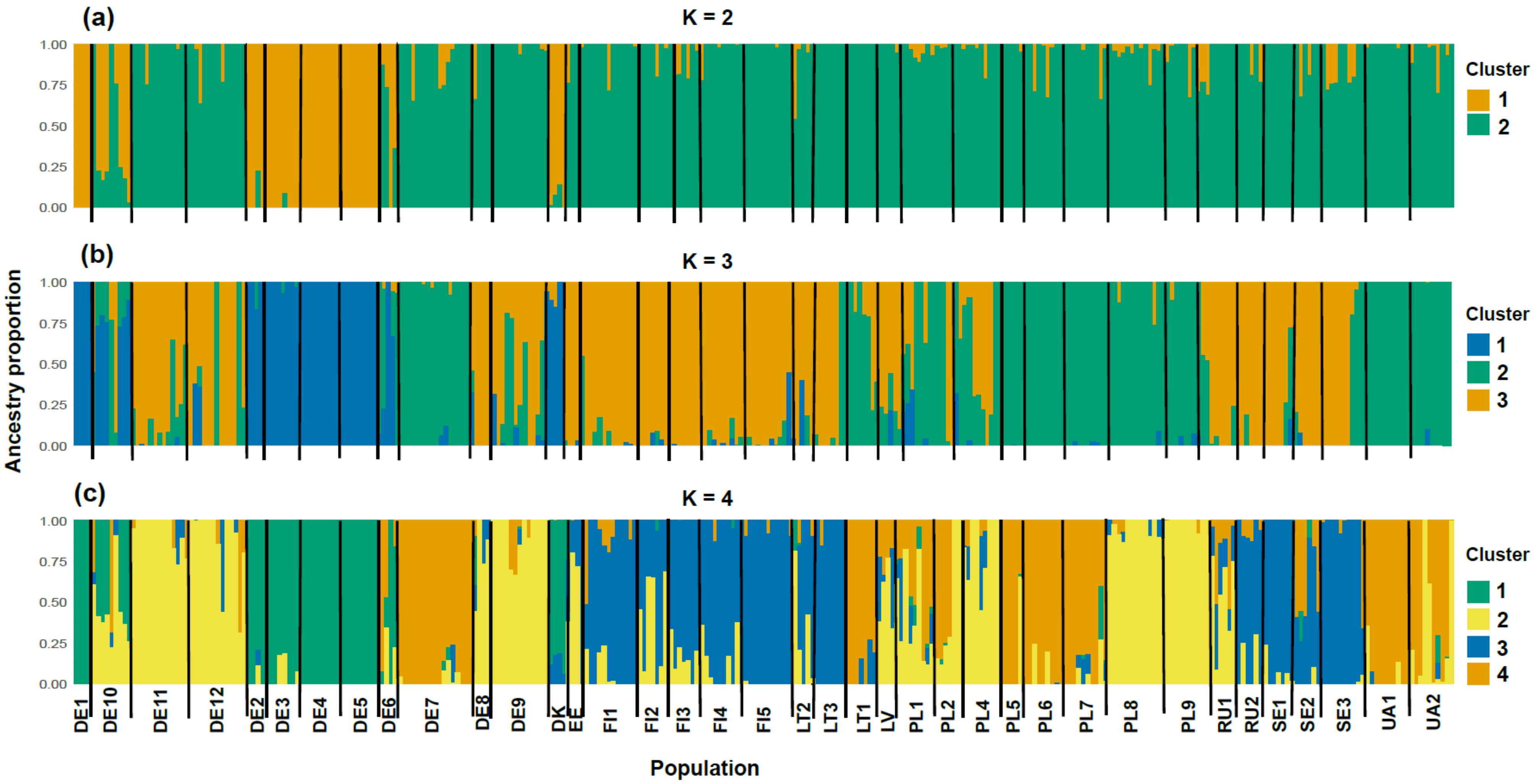


**Supplementary Figure S3.** Bar plots depicting admixture proportions calculated in ADMIXTURE for (a) two, (b) three or (c) four clusters (K). Dataset included 24,852 ddRAD-derived SNPs from 311 samples of *Drosera rotundifolia*
