## Supplementary material for "“Population genomics reveals divergent lineages across Europe in the vulnerable mire plant *Drosera rotundifolia”*": S4

**Supplementary Figure S4.** *Drosera rotundifolia* plants from six populations and three genetic clusters: DE2 - DE5 – Western cluster (W), DE7 - Eastern cluster (E) and SE1 – Northern cluster (N).

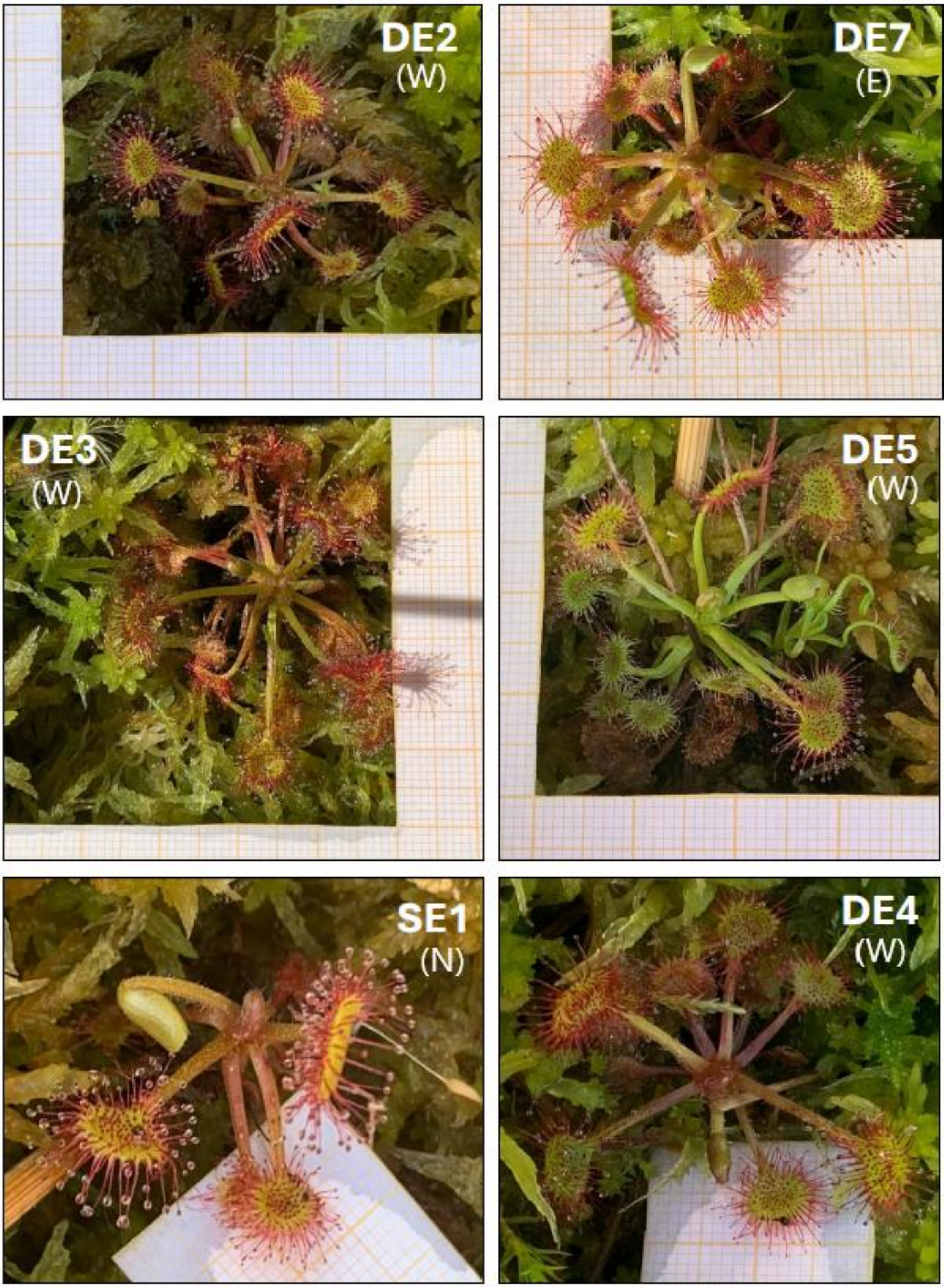
