## Supplementary material for "“Population genomics reveals divergent lineages across Europe in the vulnerable mire plant *Drosera rotundifolia”*": SF2

**Supplementary File S2.** Comparision of datasets from ipyrad pipeline with md = 6.

Selected parameters: md6, ms150, ct95

(md - minimum depth for statistical and majority rule base calling,
ms - minimum number of samples per locus for output, ct - clustering threshold)

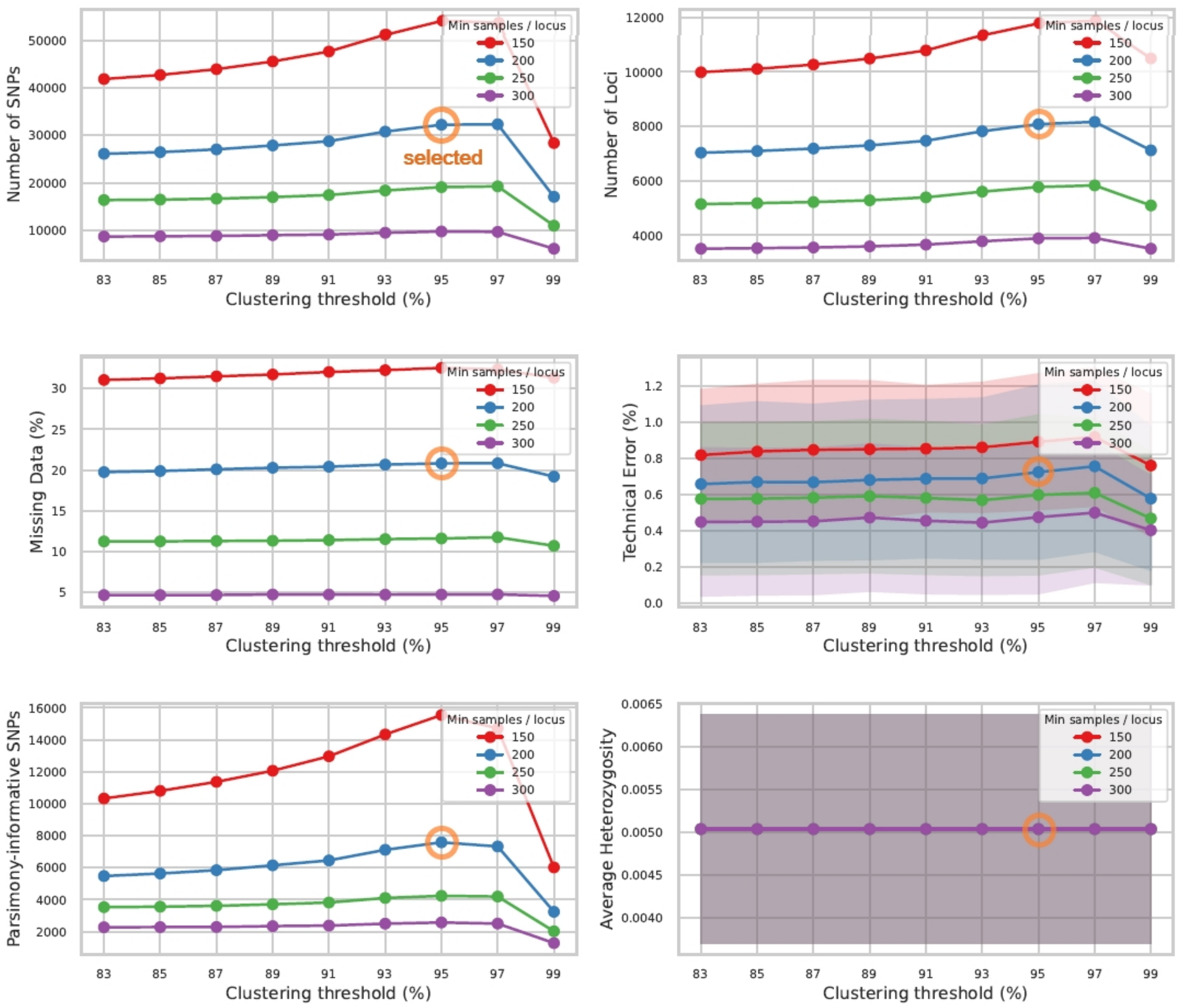

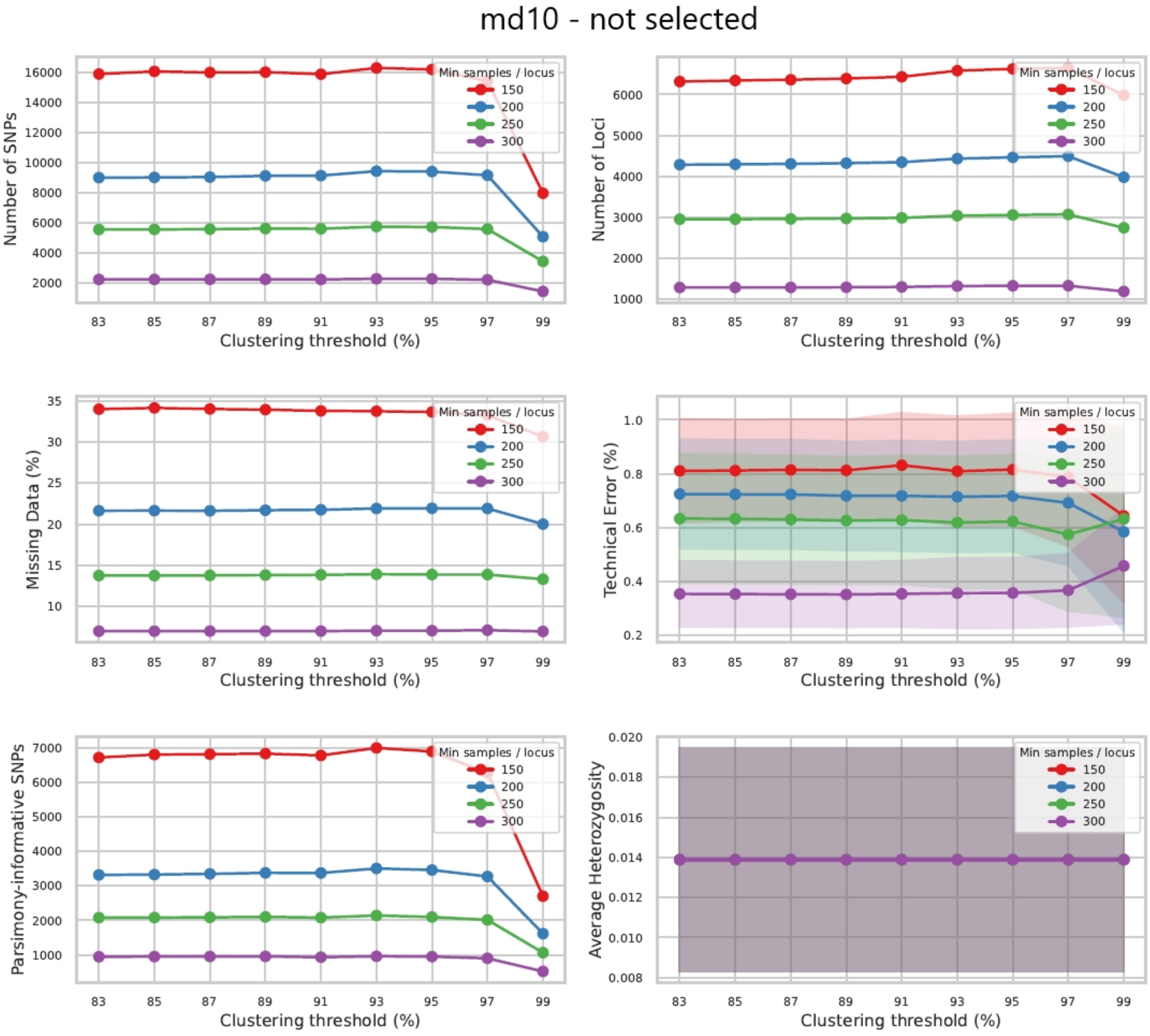

md10 - not selected
Number of SNPs
Number of Loci
Missing Data (%)
Technical Error (%)
Parsimony-informative SNPs
Average Heterozygosity
Clustering threshold (%)
Min samples / locus
150
200
250
300
