## Supplementary material for "“Population genomics reveals divergent lineages across Europe in the vulnerable mire plant *Drosera rotundifolia”*": SF3

**Supplementary File S3:** Phylogenetic tree inferred from ddRAD data for 317 samples of *Drosera rotundifolia* from 38 populations (including six pairs of technical replicates). Tree is rooted at midpoint.

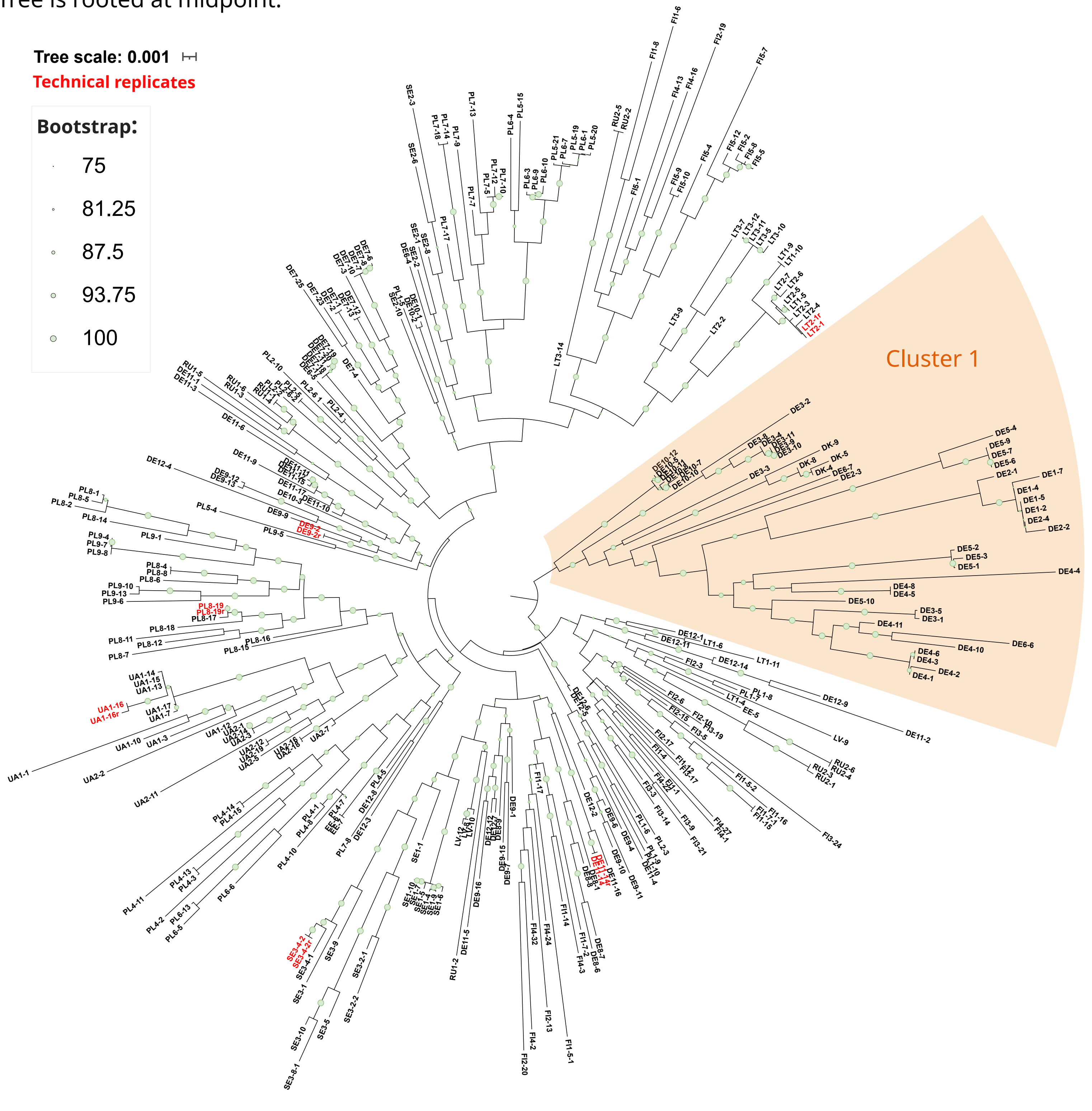
