## Supplementary material for "“Population genomics reveals divergent lineages across Europe in the vulnerable mire plant *Drosera rotundifolia”*": ST2

**Supplementary Table ST2.** Universal and cross-species primer pairs used to amplify cpDNA fragments of *Drosera rotundifolia* with their tested annealing temperatures.

| Primer name | 5'- 3' sequence | Annealing temp, °C | Publication author | Successfully sequenced samples |
|---|---|---|---|---|
| *rps*16F<br>*trnQ* | GTTGCTTTCTACCACATCG<br>GTTCGAATCCTTYCGTCCC | 59/62 | Both Saltonstall 2001 | DE10a, LV-1 |
| *trnL*"c"-F<br>*TrnL*"d"-R | CGAAATCGGTAGACGCTACG<br>GGGGATAGAGGGACTTGAAC | 52/58/59/61/ 62/63 | Both Taberlet et al.1991 | DE10a, DE1-5, FI1-1, FI5-5, LT4a, SE1-2, SE2-6, UA1-15 |
| *trnL*"e"-F<br>*trnF*"f"-R | GGTTCAAGTCCCTCTATCCC<br>ATTTGAACTGGTGACACGAG | 58/61/62 | Both Taberlet et al.1991 | DE10a, DE11-5, FI1-1, FI5-5, LT4-1, SE2-6, PL5-16, UA1-14, UA1-15 |
| *trnL-trnF*-F<br>*trnL-trnF*-R | AAAATCGTGAGGGTTCAAGTC<br>GATTTGAACTGGTGACACGAG | 59/61/62 | Both Sang et al. 1997 | SE2-6, PL5-16, UA1-14, UA1-15, FI5-5, DE1-5 |
| *atpF*<br>*atpH* | ACTCGCACACACTCCCTTTCC<br>GCTTTTATGGAAGCTTTAACAAT | 52/54/63 | Both Wang et al. 2010 | DE10a, DE1-5, FI1a, FI5-5, LT4a, UA1-15 |
| *psbK*<br>*psbI* | TTAGCATTTGTTTGGCAAG<br>AAAGTTTGAGAGTAAGCAT | 49/50/52/ 54/62 | Both Wang et al. 2010 | DE10a, LVa |
| *trnT*"a"-F<br>*trnL*"b"-R | CATTACAAATGCGATGCTCT<br>TCTACCGATTTCGCCATATC | 50/52/53/ 54/55/ 62 | Both Taberlet et al.1991 | DE7a, DE10a, DE1-5, FI1-1, FI5-5, LT4-1, SE1-2, SE2-6, PL5-16, UA1-14, LV-5, |
| aF<br>aR | ATGTCACCACAAACAGAGACTAAAGC<br>CTTCTGCTACAAATAAGAATCGATCTCTCCA | 55/62 | Both Taberlet et al.1991 | SE2-6, PL5-16, UA1-14 |
| bF<br>bR | TATCCCCTGGATTTATTTGAGGAAGGTTC<br>CGTTCGCCTTCCAATTTGCCCACTACAGT | 55/62 | Both Taberlet et al.1991 | SE2-6 |
| *RbcL*<br>*psaI* | CTAAGCCTACTAAAGGYACG<br>TGTACAAGCTCGTAACGAAGG | 58 | Both Saltonstall 2001 | No specific product obtained |
| *rpl*16F71<br>*rpl*16R662 | GCTATGCTTAGTGTGTGACTCGTTG<br>CCAACCCAATGAATCATTAGGATT | 58/59/61/ 62/63 | Jordan et al. 1996<br>Les et al. 2002 | No specific product obtained |
| *trnS*(GCU)<br>*psbD* | GCCGCTTTAGTCCACTCAGC<br>CTATGGGGTCACARCCSAGG | 62 | Hamilton 1999<br>Saltonstall 2001 | No specific product obtained |
| *trnCF*<br>*rpoB* | GRTAAAGGATTTGCAGTCCC<br>TTCTMGGGAACTCCACATACG | 59/61/62 | Both Saltonstall 2001 | No specific product obtained |
| *rps4*<br>*trnT2* | TCSTATTCCTGCAGTACAGG<br>CTGTAGGTGTAACCTTTCGC | 59/61/62/ 63 | Both Saltonstall 2001 | No specific product obtained |
| *psbA-trnH*-F<br>*psbA-trnH*-R | GTTATGCATGAACGTAATGCTC<br>CGCGCATGGTGGATTCACAAATC | 59/61/62 | Both Sang et al.1997 | No specific product obtained |
| *rps*16f<br>*rps*16r | AAACGATGTGGTARAAAGCAAC<br>AACATCWATTGCAASGATTCGATA | 52/54 | Both modified Oxelman et al.1979 | No specific product obtained |
| cF<br>cR | TGAAAACGTGAATTCCCAACCGTTTATGCG<br>GCAGCAGCTAGTTCCGGGCTCCA | 55/65 | Both Taberlet et al.1991 | No specific product obtained |
| sF<br>sR | ACTGTAGTGGGCAAATTGGAAGGCGAACG<br>GAACCTTCCTCAAATAAATCCAGGGGATA | 55/65 | Both Taberlet et al.1991 | No specific product obtained |
