## Supplementary material for "“Population genomics reveals divergent lineages across Europe in the vulnerable mire plant *Drosera rotundifolia”*": ST3

**Supplementary Table ST3.** Designed primer pairs used to amplify cpDNA fragments of *Drosera rotundifolia* with their tested annealing temperatures. Top seven primer pair were designed using reference cpDNA of *D. rotundifolia*; next five primer pairs (orange) were designed using short read approach; 12 bottom primers pairs – using long read data.

| Primer name | 5' – 3' sequence | Annealing temp, °C | Fragment size, bp | Successfully sequenced samples |
|---|---|---|---|---|
| *trnS*-F<br>*TrnG*-R | CTACGCCTTGAACCACTC<br>CTGAGGTTGTTGGAATTATTCG | 55 | 906 | SE2-2, PL8-1, UA1-15 |
| *PsbI*-F<br>*ycf15*-R | TCTCAAGACAGTCAAGCC<br>AAGAAAGGACTCACTGAG | 50 | 920 | SE2-2, PL8-1, UA1-15 |
| *psbM*_b-F<br>*tRNA*_A-R | AGGCAGTAGGAACTAGAATG<br>CTGTCAAGGCGGAAGTTG | 50 | 707 | SE2-2, PL8-1, UA1-15 |
| *rps4*_b-F<br>*NdhB*-R | GACTGAGAAAATGGCAGAG<br>TTAGGCGATGTATTCTGG | 50 | 485 | SE2-2, PL8-1, UA1-15 |
| *tRNA*_Asp-F<br>*tRNA*_Tyr-R | ATTCTATGTCTCCCTTTCG<br>CTACGCTGGTTCAAATCC | 50 | 518 | SE2-2, PL8-1, UA1-15 |
| *petB*-F<br>*petB*-R | GTTCTAATATGAATCTGAGG<br>AATTTCATCTCATACAGCTC | 55 | 314 | SE2-2, PL8-1, UA1-15 |
| *atpB*<br>*rbcL* | ACATCCAGTACCGGACCAATAA<br>AACACCCGCTTTGAATCCAA | 55 | 288 | SE2-2, PL8-1, UA1-15 |
| 329-F<br>757-R | GGACACAACTTGAACATTC<br>GACTAATCTTCTTTGCTCC | 52/54 | 444 | SE2-2, LV-3, LT3-2, FI1-1, FI5-5, DE1-5, UA1-5, DK-8 |
| 101524-F<br>102199-R | GGAACTATCAAGACAACAC<br>GAATACCCAGAGCAATTTC | 52/54 | 684 | SE2-2, LV-3, LT3-2, FI1-1, DE1-5, DK-8 |
| 130299-F<br>130694-R | GACTAAGGTGAACGAACC<br>GTACCATAGGAAGGAATG | 52/54 | 368 | SE2-2, LV-3, LT3-2, FI1-1, FI5-5, DE1-5, DK1-8 |
| 174151-F<br>174619-R | CCTCCTTGATCCTATCCATC<br>GTAGACAAGACTCGCATC | 52/54 | 469 | SE2-2, LV-3, LT3-2, FI1-1, FI5-5, DE1-5, UA1-15, DK-8 |
| 180053-F<br>180717-R | CGTTATTGTTGCTGTGATC<br>GAGCGTCCTTTCCTATTCC | 52/54 | 681 | SE2-2, LV-3, LT3-2, FI1-1, DE1-5, DE7-2, DE4-1 |
| *rps14*/*psaB*-F<br>*rps14*/*psaB*-R | GTTCTTTCTTTGAGGATCG<br>GGGATTTGGATTCTATGG | 54 | 491 | DE7-1, DE1-5, SE3-6, UA1-15, FI1-1, FI5-5, PL8-1, LT1-4 |
| A3229-gt022/*psbJ*-F<br>A3229-gt022/*psbJ*-R | GGAACTATCAAGACAACAC<br>CCATCTTTCGTACCTCTTC | 58 | 838 | DE7-1, DE1-5, SE2-6, UA1-15, FI5-5 |
| 3229-gr031/ps15-F<br>3229-gr031/ps15-R | CGGATCTTTCTCCTGCTC<br>TCCTCTGTTGATGTTGAG | 54 | 652 | DE7-1, DE1-5, DE11-5, SE6-6, UA1-15, FI5-5 |
| 3229-gt031-F<br>3229-gt031-R | GGTGACCAATTAACTAACC<br>AGGGAAACTCGCTCTTGC | 54 | 723 | DE7-1, DE1-5, SE2-6, UA1-15, FI5-5 |
| 33049-F<br>33438-R | CGTTACAAAGAAGTTCAGG<br>CAAGAGGGATGTGGTTGT | 54 | 426 | DE7-1, DE1-5, SE2-6, UA1-15, FI5-5 |
| 19051-F<br>19729-R | ATTATATGTTTCGGCGGG<br>TTCATATCGTCAACTCTGC | 54 | 714 | DE7-1, DE1-5, SE2-6, UA1-15, FI5-5 |
| 60985-F<br>61870-R | GAATAGAATGAGATTTCCGC<br>AGCAACTAGGCTCTCTAC | 54 | 548 | DE7-1, DE1-5, SE2-6, UA1-15, PL8-1, LT1-4, DK-1 |
| 74467-F<br>75402-R | ACTGGGACTGAGACAAGG<br>CGCATTTCACTGCTACAC | 58 | 363 | DE7-1, SE2-6, UA1-15, FI5-5, DK-8 |
| PsaC-F<br>PsaC-R | CCATATTACTGTACCTTGTAGC<br>CTTGATGAGGGGATTTGT | 54 | 922 | DE7-1, DE1-5, FI5-5, SE2-6, UA1-15 |
| 188668-F<br>189113-R | GTACCTTACATTTCAGATCC<br>AAGTTCTTCAGTAGGGTC | 54 | 482 | DE7-1, SE2-6, UA1-15, FI5-5 |
| 116633-F<br>117576-R | CTAAACCACTAGACGATGG<br>TGAACCGATGACTTACGC | 54 | 979 | DE7-1, DE1-5, SE2-6, UA1-15 |
| 141823-F<br>142395-R | GGATCTATGCTTCTACTC<br>AATCGGTATTGGAACTGC | 54 | 607 | DE2-1, DE1-5, SE2-6, UA1-15, FI5-5 |
